# Quinazolinone and Phthalazinone Inhibitors of the HDAC6/Ubiquitin Protein-Protein Interaction

**DOI:** 10.64898/2025.12.18.695271

**Authors:** Sydney Gordon, Jordi C.J. Hintzen, Sebastian Dilones, Brockton Keen, Callie E.W. Crawford, George M. Burslem

**Affiliations:** Department of Biochemistry and Biophysics, University of Pennsylvania, PA 19104; Department of Cancer Biology, University of Pennsylvania, PA 19104; Epigenetics Institute, Perelman School of Medicine, University of Pennsylvania, PA 19104

**Keywords:** HDAC6, PPI Inhibitor, Protein-Protein Interactions, Zinc Finger Ubiquitin Binding

## Abstract

Histone deacetylase 6 (HDAC6) is a class IIb histone deacetylase that regulates diverse cytosolic acetylation through its two catalytic deacetylase domains and a C-terminal zinc finger ubiquitin-binding domain (ZnF-UBD). This ZnF-UBD mediates key protein–protein interactions (PPIs) that couple deacetylation and ubiquitin-dependent degradation. While most HDAC6 inhibitors target the catalytic domains, the ZnF-UBD represents an underexplored target. Here, we validate previously reported small-molecule inhibitors of the HDAC6 ZnF-UBD/ubiquitin interaction and describe novel N-alkyl moieties based on quinazolinone and phthalazinone scaffolds. Starting from known quinazolinone and phthalazinone scaffolds, a literature and modeling-guided scaffold hop revealed potential for an extended phthalazinone series. Results obtained both in fluorescence polarization (FP) and differential scanning fluorimetry (DSF) confirm this hypothesis. Additionally, late-stage diversification yields compounds with improved predicted physicochemical properties. Finally, machine-learning-based co-folding affinity predictions correlate with experimental IC□□ rank order, highlighting their utility in PPI inhibitor design. These studies continue expanding the chemical space of HDAC6 ZnF-UBD inhibitors and build upon existing foundations for future therapeutic and mechanistic exploration of HDAC6– ubiquitin signaling.

## Introduction

Histone deacetylase (HDAC) proteins are a class of enzymes associated with the deacetylation of lysine.^1^ Removal of acetyl groups from histone tails decreases gene expression by reducing charge separation between DNA and positively charged lysine residues.^2^ However, some HDACs play roles beyond the nucleus, including catalyzing the removal of acetyl groups from non-histone substrates,^3, 4^ catalyzing other chemical reactions ^5-10^, and scaffolding protein complexes.^11^

Across the 18 human HDACs, histone deacetylase 6 (HDAC6) is unique in its structural and functional properties.^2^ HDAC6 is primarily cytosolic, with a broad topography of substrates including cortactin,^12^ Hsp90 (heat shock protein),^13^ a-tubulin,^14^ α-catenin,^15^ ERK1^16^ and FoxP1^17^. Additionally, HDAC6 is the only HDAC that contains two deacetylase domains (DAC1 and DAC2) as well as a C-terminal zinc finger-ubiquitin binding domain (ZnF-UBD).^18^ The DAC1 and DAC2 domains are responsible for facilitating the catalytic deacetylase activity of the protein. However recent literature suggests that the DAC1 and DAC2 domains may also function both in protein lactylation^5, 6^ and ubiquitination^7, 9^. DAC1 is hypothesized to facilitate or directly perform the transfer of ubiquitin onto the autophagy regulatory protein ATG3^7^ and the DNA repair protein, MSH2, following deacetylation of K845, K847, K871, and K892.^9^

Interestingly, the ZnF-UBD domain also plays an important role in protein degradation pathways.^19^ While the ZnF-UBD is non-catalytic, it mediates autophagy, mitophagy, and stress granule clearance through binding to polyubiquitinated substrates and shuttling misfolded complexes and damaged mitochondria to the budding aggresome.^8, 20, 21^ This UBD polyubiquitin binding activity is primarily facilitated through the R1155 and Y1184 residues at the UBD site. Abolishing these residues leads to loss of ZnF-UBD ubiquitin binding.^22, 23^ Additionally, the mechanism for ZnF-UBD recognition of polyubiquitinated substrates has yet to be fully realized. Some research suggests that the UBD can recognize K63 linked polyubiquitin chains specifically,^7^ while other literature suggests that instead of ubiquitin linkage recognition, the ZnF-UBD specifically binds to unconjugated ubiquitin produced by ataxin-3 cleavage at protein aggregate sites.^19^ Regardless, ZnF-UBD plays an integral role in the lysosomal degradation pathway through ubiquitin binding. In a disease-relevant context, the ZnF-UBD is known to interact with a variety of pathologies including breast cancer, colorectal cancer, chronic inflammation, viral infection, Alzheimer’s disease, and multiple myeloma, making it a promising therapeutic target.^8, 24^ Given this role in pathogenesis, and our interests at the intersection of acetylation and degradation,^4^ we sought to develop inhibitors of the HDAC6 ZnF-UBD/Ubiquitin protein-protein interaction.

## Results and Discussion

Previous crystallographic studies of the HDAC6 ZnF-UBD in complex with both ubiquitin,^19^ ubiquitin derived peptides, and fragments provided rational starting points for inhibitor development.^25, 26^ We initially synthesized a previously reported ZnF-UBD inhibitor, Compound 31, bearing an alkyl carboxylate chain that mimics the diglycine motif of ubiquitin and a quinazolinone core that forms Π-Π interactions with W1182 and Y1184 in the ZnF-UBD (Figure 1 A/B).^25^ We were able to recapitulate fluorescence polarization assays to show that Compound 31 is able to compete with a fluorescently-tagged peptide mimicking the ubiquitin C-terminus with an IC_50_ of 3.1 ± 1.2 µM (Fig. 1B/C). Gratifyingly, this IC_50_ value was similar to the literature value for Compound 31 (2.3± 0.6 µM).^25^ These results provided a useful starting point for further compound elaboration.

**Figure 1:**
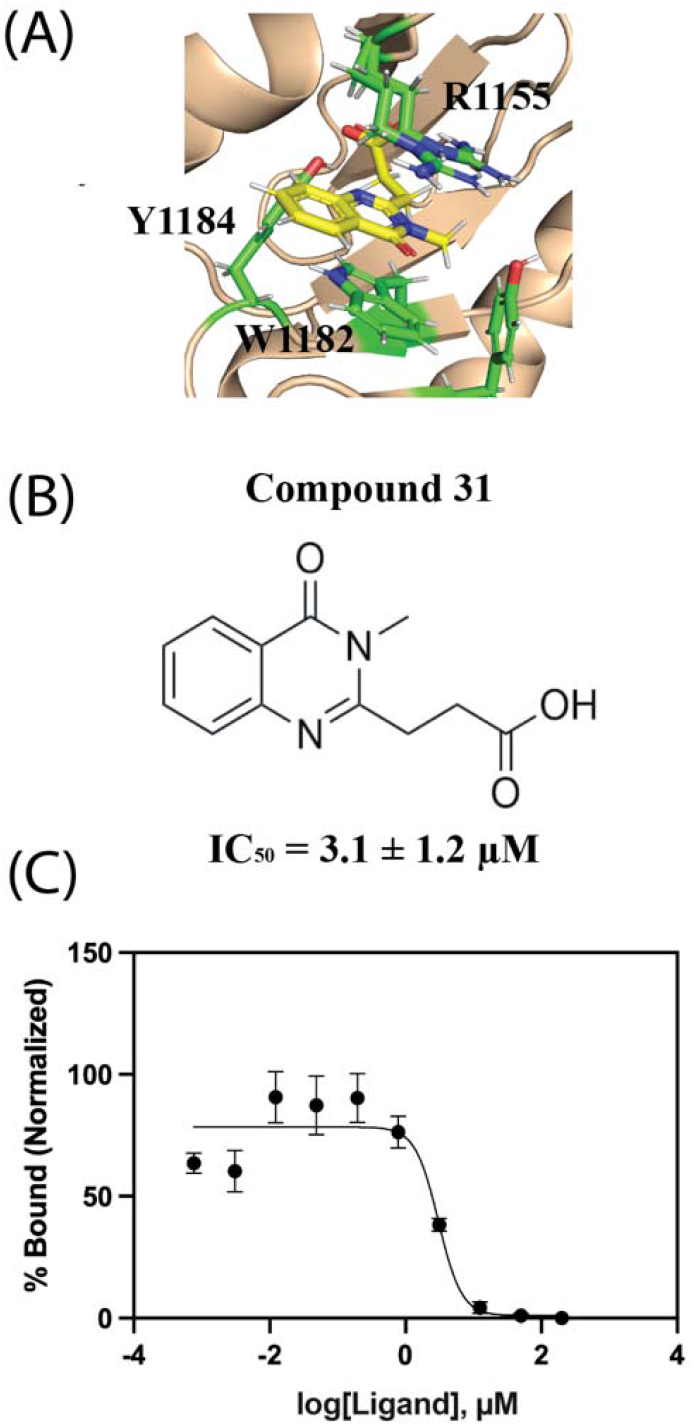
Compound 31 Inhibits the HDAC6 ZnF-UBD. (A) Molecular structure of Compound 31 bound to the HDAC6 ZnF-UBD from PDB ID: 6CED. (B) Structure of Compound 31 and IC_50_ with error reported as SEM. (C) Fluorescence polarization competition assay binding curve measuring Ub RLRGG-FITC peptide displacement from the ZnF-UBD by Compound 31.

We first verified the importance of the *N*-methyl group in Compound 31 and the length of the alkyl spacer between the quinazolinone core and the carboxylic acid^25, 27^. It has been highlighted previously that replacement of the carboxylate or extension of the alkyl chain results in total loss of compound binding (ref. 25, Compound 7, Compound 25-27)^25 27^. We similarly find that modifying the length of the alkyl acid chain results in significant loss of activity and confirm that loss of the *N*-methyl has the same result, thus reinforcing the importance of the ethyl carboxylate chain in stabilizing the ligand at the UBD binding site (Table 1).

**Table 1.**
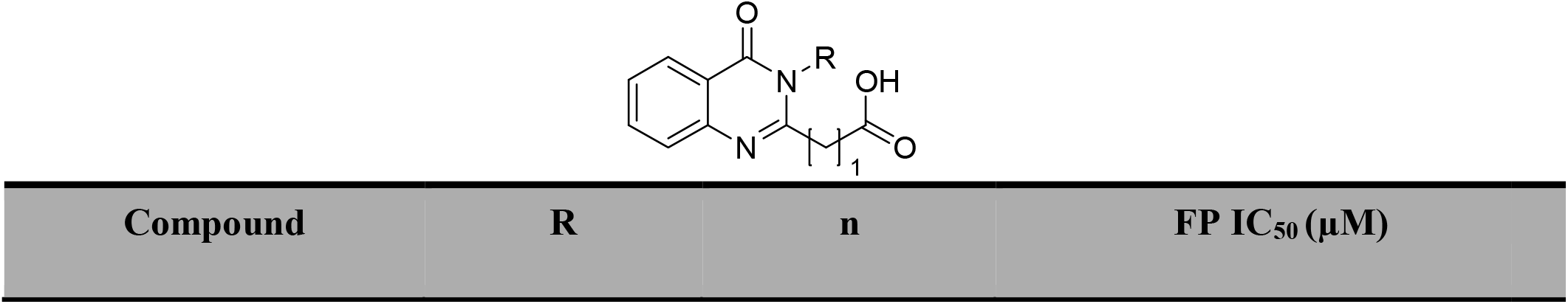
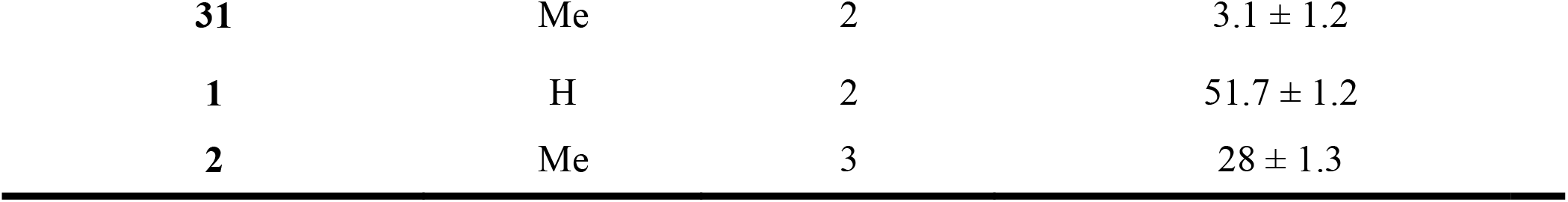
Initial Structure Activity Relationships.

Informed by molecular modeling and analysis of existing crystal structures which show a accessible pocket in HDAC6, proximal to the C-terminal Ub binding site,^26^ we sought to expand compounds via derivatization of the quinazolinone core along the methyl vector. Early exploration and literature demonstrated benzyl and benzyl derivatives were tolerated in place of *N*-methylation (ref. 27, Compound 29, Compound 27, Compound 32), and these findings were confirmed through FP competition assays (Table 2). In parallel to our studies, the structural genomics consortium reported on further derivatized compounds, including SGC-UBD253, informing our choice of side chain in Compound 6.^27^

**Table 2.**
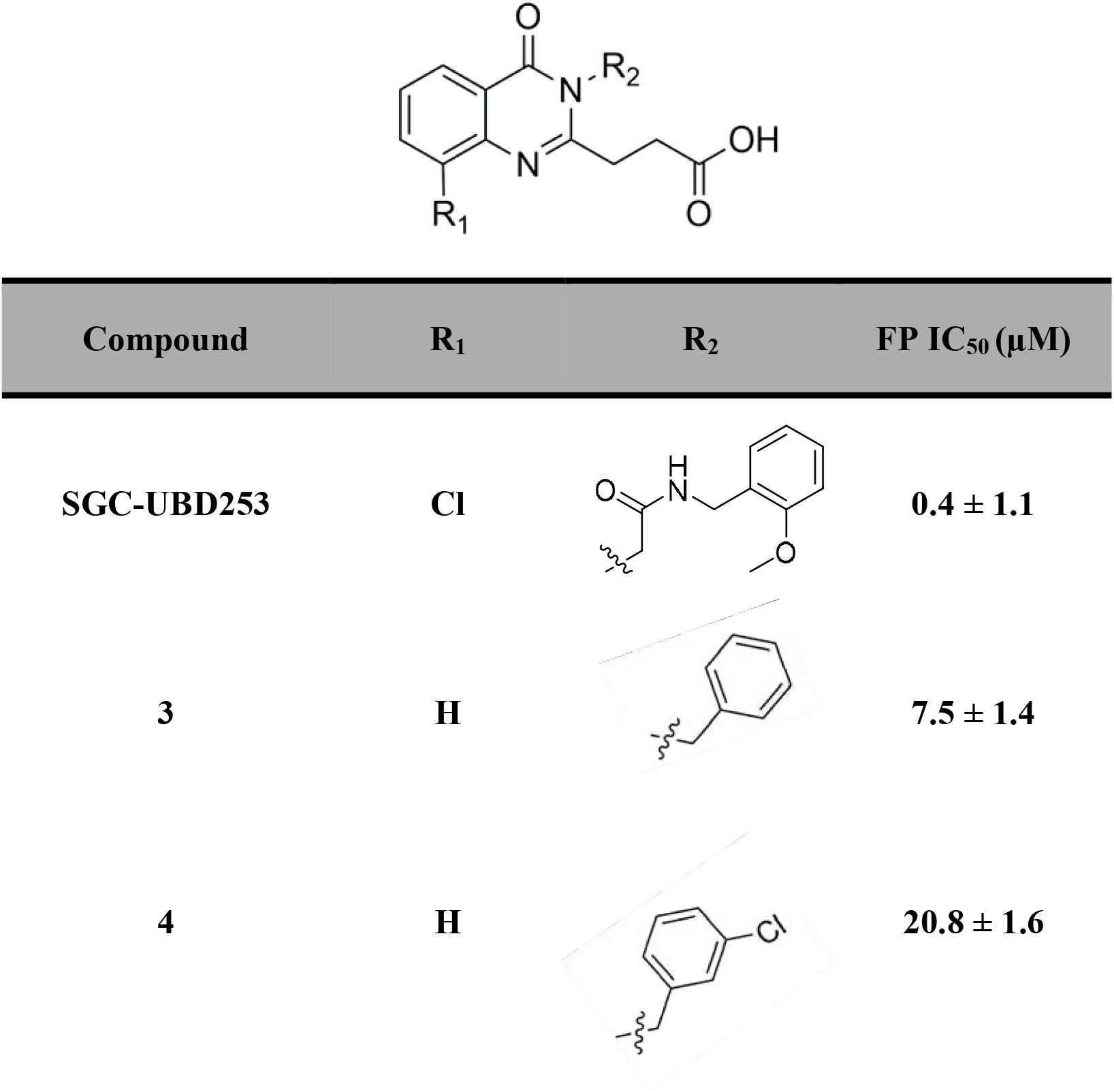
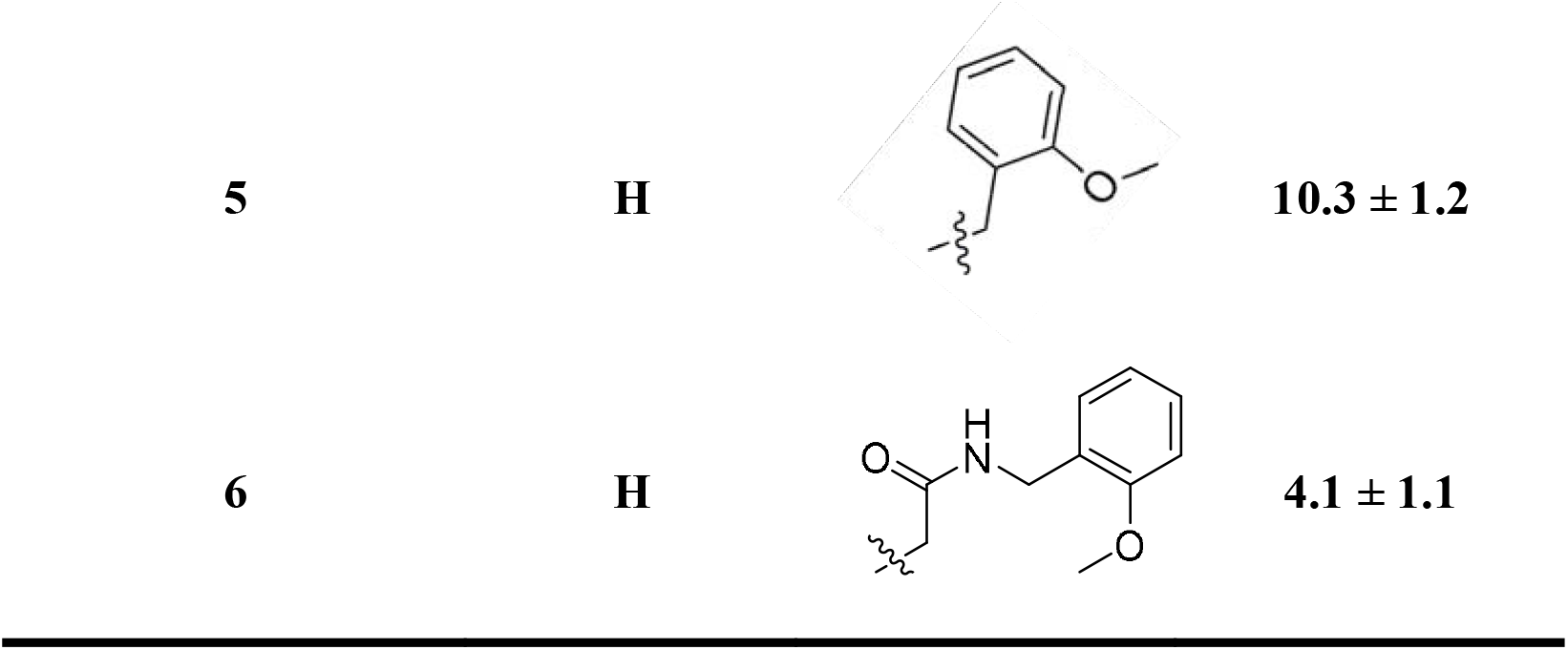
*N*-Alkyl Quinazolinone Derivatives.

### Expanded Sidechain Library

Utilizing SGC-UBD253 as a positive control^27^ we explored additional amide sidechains, seeking to retain activity but improve drug-like properties as scored by our recently reported chemo-informatics tool, LOSERS, for lead discovery.^28^ Virtual library enumeration using libraries of commercially available amines, followed by chemoinformatic scoring and docking against the HDAC6 binding site led us to prioritize the synthesis of several derivatives (Figure 2), considering both improved physicochemical properties and predicted binding modes.

**Figure 2:**
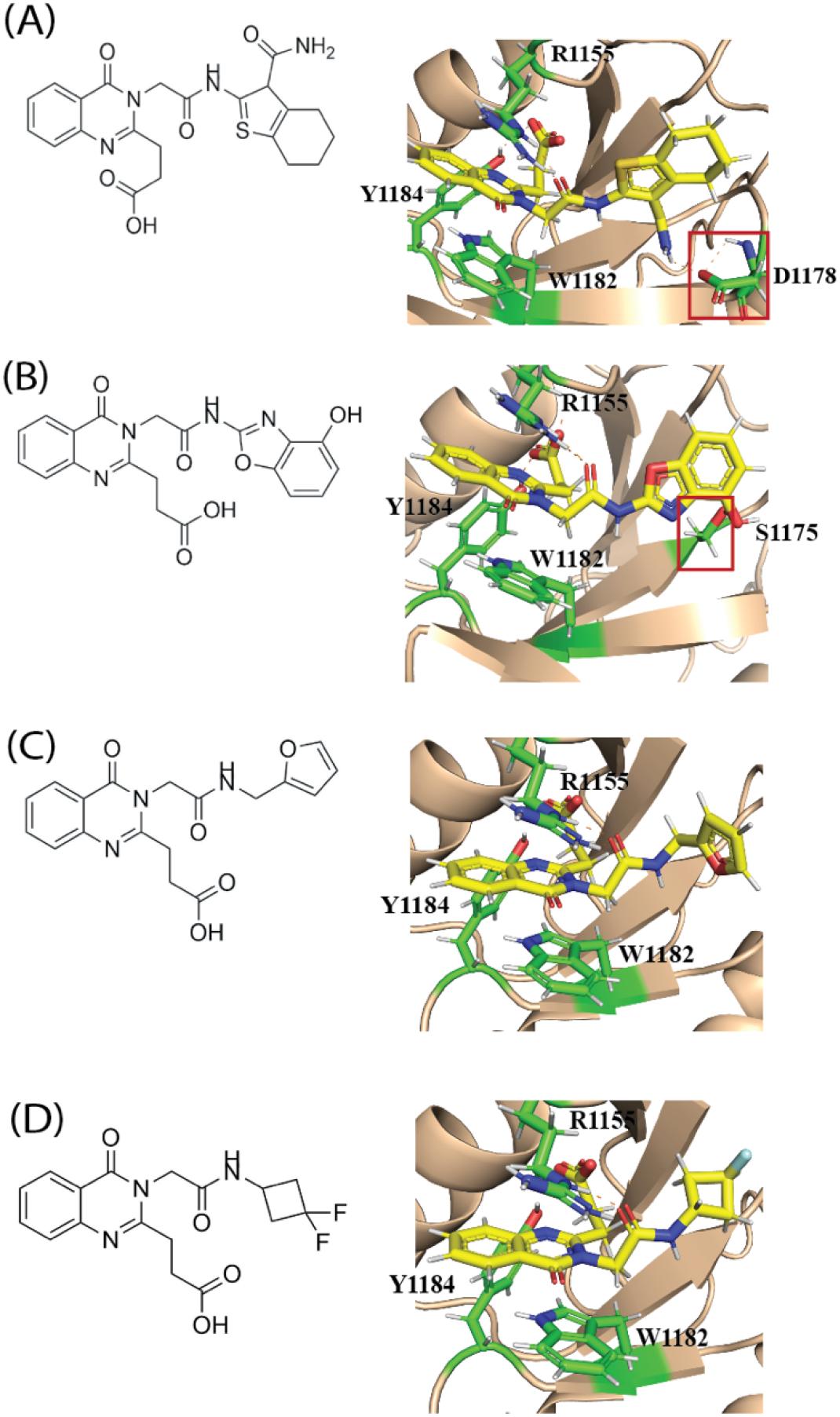
Docking Structures of expanded N-alkyl quinazolinone derivatives. Structures and docked poses of (A) Compound 7, (B) Compound 8, (C) Compound 9, (D) Compound 10 into the HDAC6 ZnF UBD.

The 2-methoxybenzylamine could be readily substituted with several other heterocycles without substantial impact, although the benzoxazole substitution was not tolerated (Fig. 2B, Table 3). Based on the postulated binding mode, Compound 7 may pick up an additional polar interaction in the binding site at D1178 (Fig. 2A). Both Compound 9 and Compound 10 (Fig. 2C/D, Table 3) demonstrate comparable ZnF-UBD affinity and Compound 10 has a significantly improved lead likeness (LOSERS Score of 73.5 compared to 46.9 for SGC-UBD253)^28^, due to reduced lipophilicity and number of aromatic rings (See Table S1 for all calculated properties used for LOSERS Scoring). We postulate that these compounds could also be further improved by the addition of a chloride at the 8-position, analogous to SGC-UBD253 (Table 2/3).

**Table 3.** Expanded *N*-Alkyl Quinazolinone Derivatives.

| Compound | R | FP IC<sub>50</sub> (μM) |
| --- | --- | --- |
| 7 |  | 4.0 ± 1.3 |
| 8 |  | > 200 |
| 9 |  | 4.9 ± 1.7 |
| 10 |  | 2.4 ± 1.3 |

### Phthalazinone Scaffold

In parallel to our development of the quinazolinone series above, we sought to investigate alternative heterocyclic cores. Although a phthalazinone analogue has been reported previously, robust inhibition of the ZnF-UBD/ubiquitin interaction had not been demonstrated (ref. 25, Compound 19). We sought to determine if a quinazolinone to phthalazinone scaffold hop could be tolerated. Guided by molecular docking and our success within the expanded quinazolinone series (Table 3), we developed a virtual library focused on developing N-alkyl linkers of the phthalazinone (Fig. 3A/B, Table 4). Bolstered by docking poses, we synthesized a small library of phthalazinone derivatives which revealed that the scaffold hop was tolerated, resulting in the identification of several new analogues. Compound 12, which is the closest in structural composition to Compound 6 retains similar activity, including an identical linker between the heterocyclic core and acid sidechain. Similarly, N-alkyl derivatives of the quinazolinone scaffold could be supplanted onto the phthalazinone core, suggesting a similar binding mode consistent with the docking poses (Figure 3).

**Table 4.**
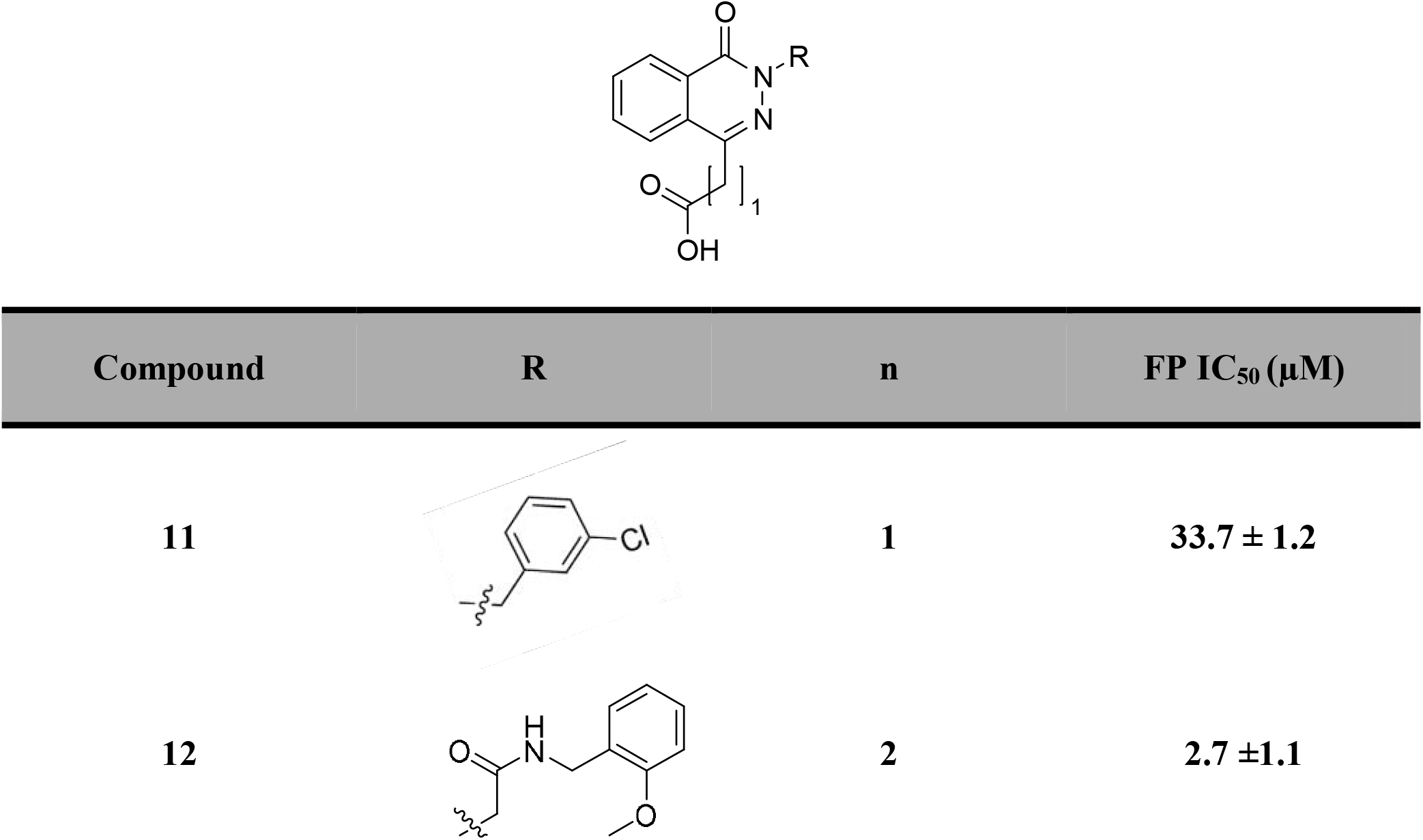
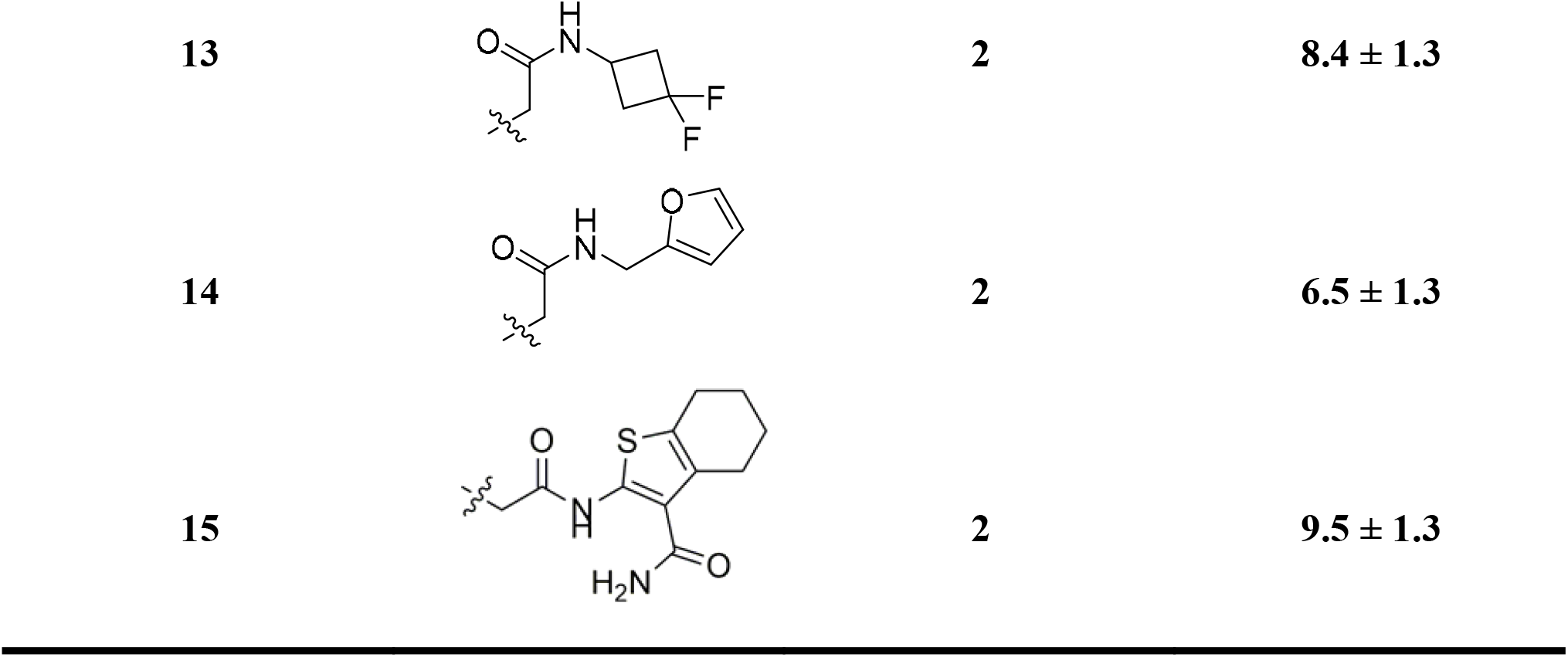
Phthalazinone Derivatives.

| Compound | R | n | FP IC<sub>50</sub> (μM) |
| --- | --- | --- | --- |
| 11 |  | 1 | 33.7 ± 1.2 |
| 12 |  | 2 | 2.7 ± 1.1 |
| 13 | | 2 | $8.4 \pm 1.3$ |
| 14 | | 2 | $6.5 \pm 1.3$ |
| 15 | | 2 | $9.5 \pm 1.3$ |

**Figure 3.**
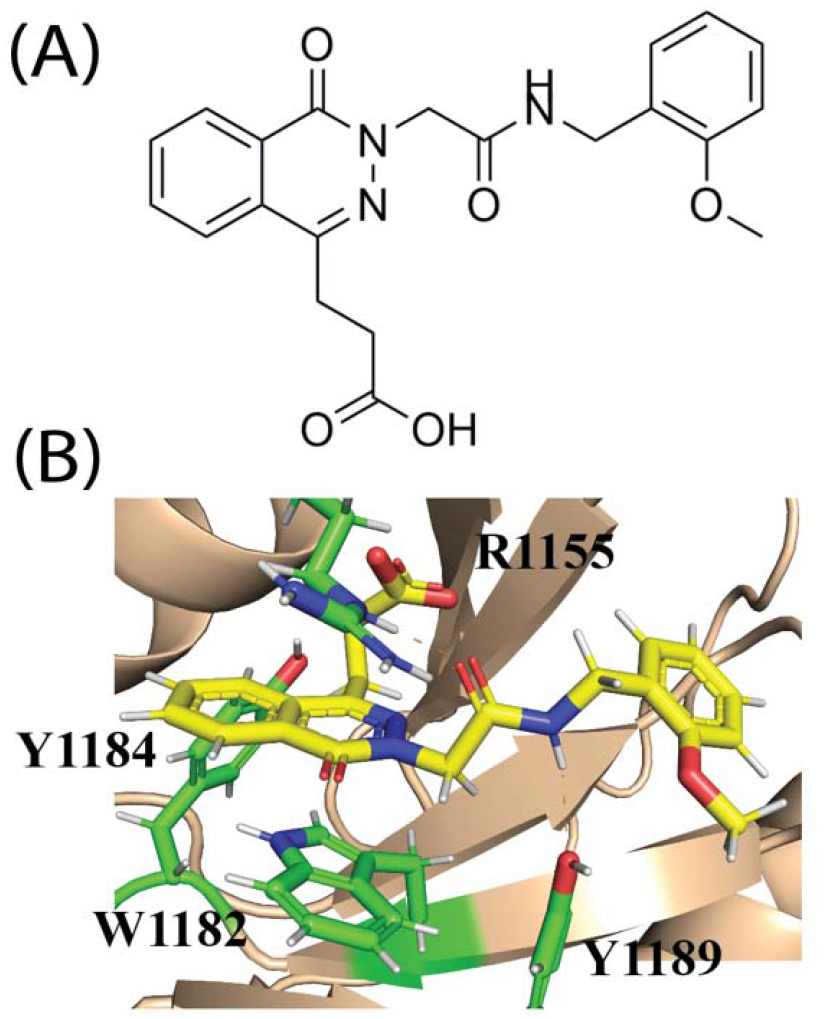
Molecular Modeling of Compound 12. (A) Structure of Compound 12. (B) Predicted binding pose of Compound 12 in the HDAC6 ZnF-UBD.

### Assessing Thermal Stability of Lead Compounds

Following FP analysis, we selected lead compounds (Compound 7, Compound 10, Compound 12) for further biochemical assessment. Using DSF, we were able to measure the thermal stability of purified ZnF-UBD in the presence of our three lead compounds (Figure 4B). Use of the validated ZnF-UBD inhibitor, SGC-UBD 253 functioned as an essential positive control for these experiments (Figure 4A).^27^ Gratifyingly, treatment with Compounds 7, 10 and 12 exert a stabilizing effect on the ZnF-UBD. Amongst these compounds, the phthalazinone Compound 12 exerts a 6.9 C° melting temperature shift, further validating the tolerance of a quinazolinone to phthalazinone scaffold hop (Figure 4B).

**Figure 4:**
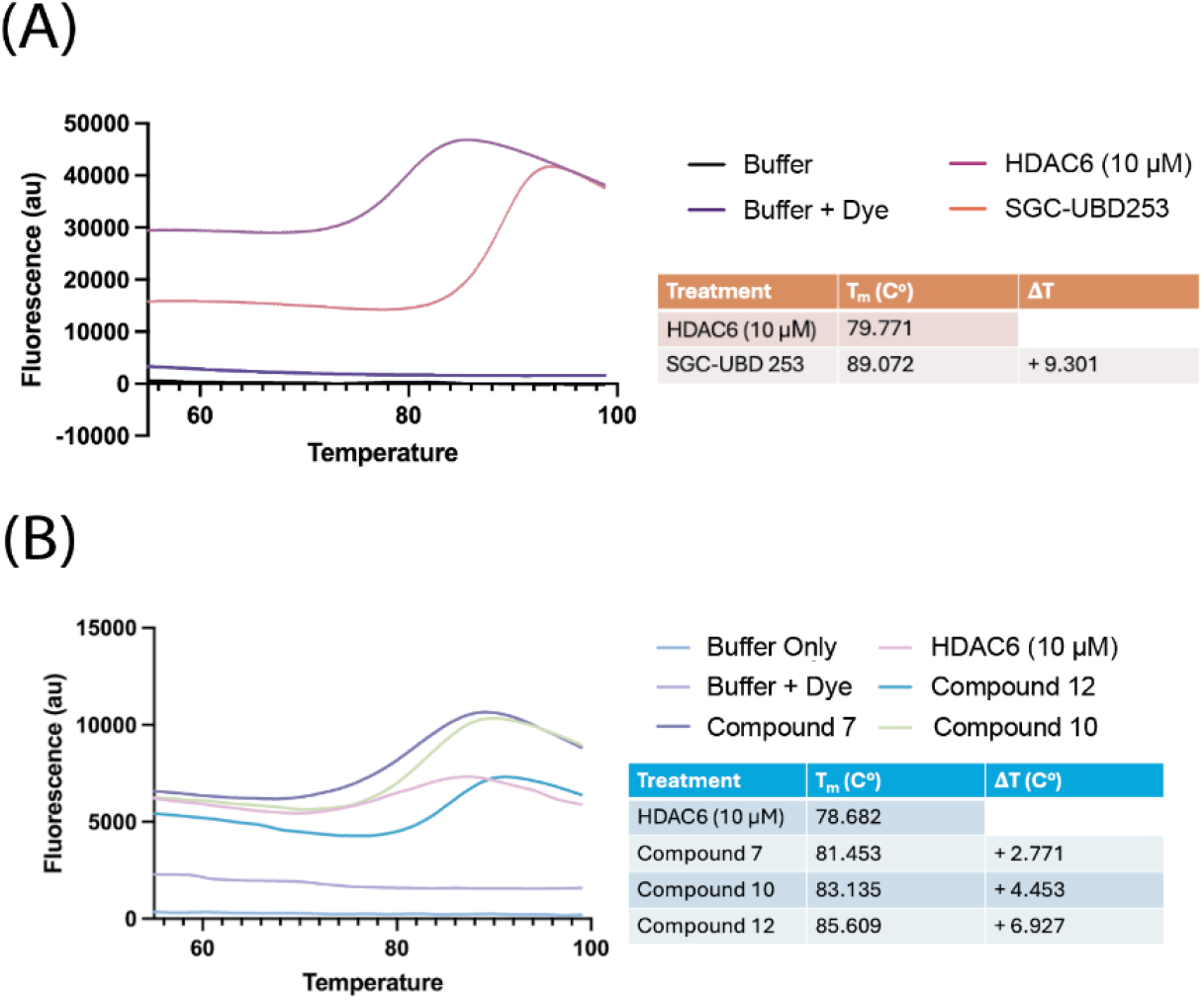
Differential scanning fluorimetry (DSF) results of SGC-UBD253, Compound 7, Compound 10, and Compound 12. (A) Shift in ZnF-UBD melting temperature (T_m_) upon treatment with 50 µM SGC-UBD253 positive control. All conditions were repeated in technical triplicate. (B) Shift in ZnF-UBD melting temperature (T_m_) upon treatment with 50 µM Compound 7, Compound 10, and Compound 12. All conditions were repeated in technical triplicate.

### Comparison of Co-Folding to Docking

Finally, we sought to retrospectively compare the use of co-folding ligand scoring methods to traditional molecular docking in guiding PPI inhibitor design. Taking a subset of compounds with a range of IC_50_ values from our studies, we performed co-folding analysis and IC_50_ predictions using Boltz-2 (Table 5).^29^ Boltz-2 predictions were able to successfully identify SGC-UBD253 as the best compound and other predicted IC_50_ values and subsequent ranks showed reasonable correlation with experimental results. However, it outperformed docking-based ranking in this compound series (Figure 5), suggesting it may be more robust and reliable moving forward than docking for the development of inhibitors of protein-protein interactions.

**Table 5.**
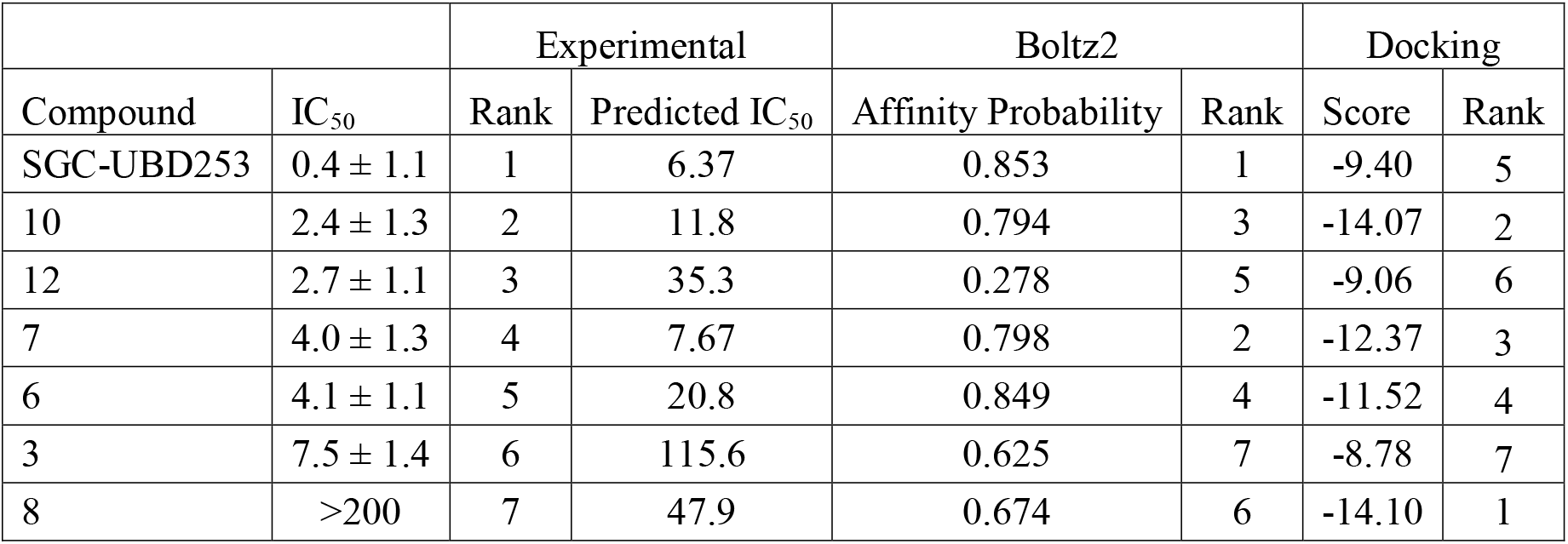
Comparison of Experimental and Predicted IC_50_ Values and Compound Ranking.

| Compound | $IC_{50}$ | Experimental | | Boltz2 | | Docking | |
| --- | --- | --- | --- | --- | --- | --- | --- |
| | | Rank | Predicted $IC_{50}$ | Affinity Probability | Rank | Score | Rank |
| SGC-UBD253 | $0.4 \pm 1.1$ | 1 | 6.37 | 0.853 | 1 | -9.40 | 5 |
| 10 | $2.4 \pm 1.3$ | 2 | 11.8 | 0.794 | 3 | -14.07 | 2 |
| 12 | $2.7 \pm 1.1$ | 3 | 35.3 | 0.278 | 5 | -9.06 | 6 |
| 7 | $4.0 \pm 1.3$ | 4 | 7.67 | 0.798 | 2 | -12.37 | 3 |
| 6 | $4.1 \pm 1.1$ | 5 | 20.8 | 0.849 | 4 | -11.52 | 4 |
| 3 | $7.5 \pm 1.4$ | 6 | 115.6 | 0.625 | 7 | -8.78 | 7 |
| 8 | >200 | 7 | 47.9 | 0.674 | 6 | -14.10 | 1 |

**Figure 5:**
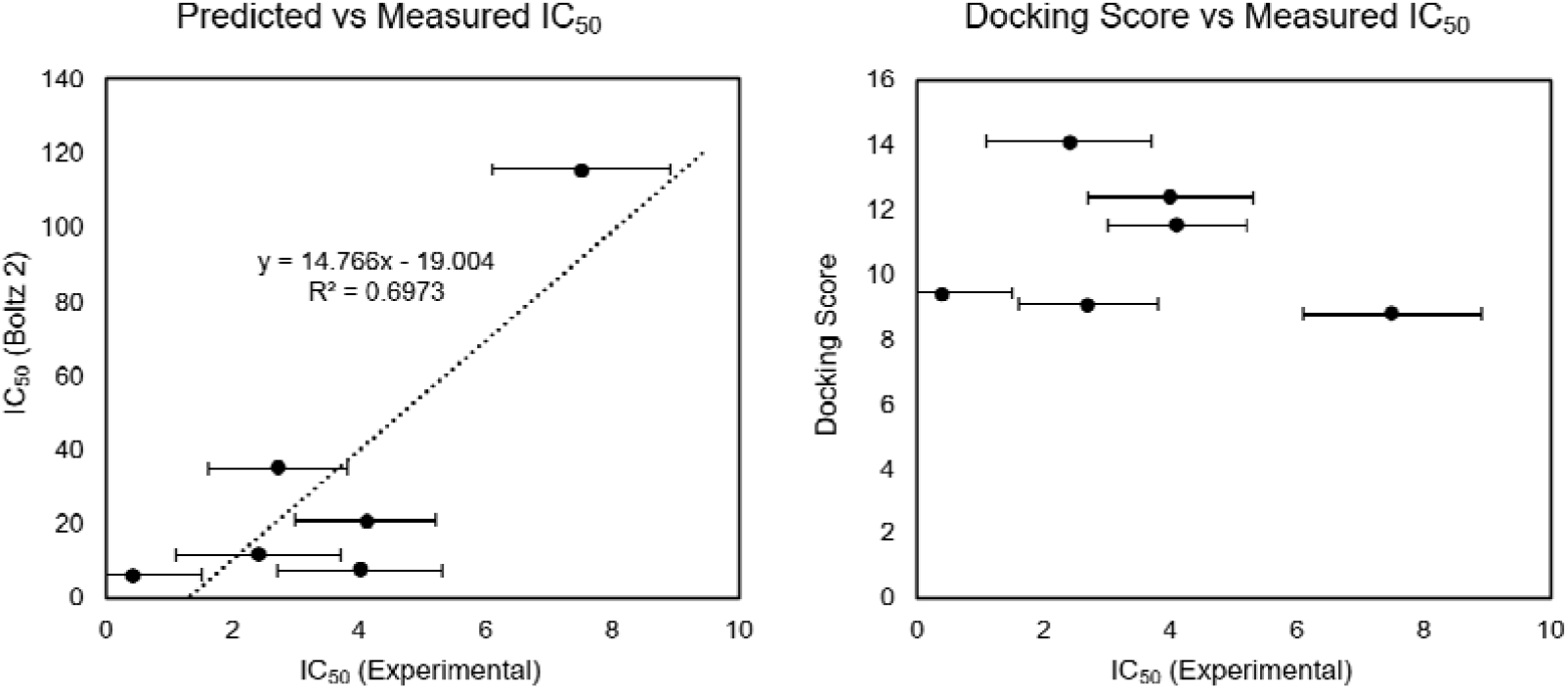
Comparison of Co-folding and Docking. (Left) Boltz2 was used to predict IC_50_ values and compared to experimental results for all predictions. (Right) Docking scores were compared to experimental results.

## Discussion

Here, we identify new derivatives, Compound 10 and Compound 12, which inhibit the ZnF-UBD/Ub interaction in FP competition assays (2.4 ± 1.3 µM and 2.7 ± 1.1 µM respectively) and demonstrate substantial thermal stability shifts in DSF. Compound 10 shows increased lead/drug likeness according to its calculated properties, which makes it an interesting candidate for further development as a chemical probe for cell-based exploration and potential future *in vivo* studies. Furthermore, while a single phthalazinone compound has been reported in the ZnF-UBD literature previously, we demonstrate that altering the exit vector of the ethyl carboxylate and optimizing substituents can result in compounds with equivalent potency to the quinazolinone series, strongly suggesting a conserved binding mode. Future work will focus on continued expansion of the phthalazinone library through developing additional N-alkyl substitutions and exploring modifications at additional positions on the core. We will report cell based characterization of our further optimized derivatives in due course.

Finally, we demonstrate that predictive co-folding models hold potential for the design and optimization of inhibitors of protein-protein interactions. Low affinity probability scores for the phthalazinone series, could in part be explained by limited examples of phthalazinones in the medicinal chemistry literature, and therefore the training data available. In conclusion, we revalidate and develop upon the chemical space of inhibitors of the HDAC6 ZnF-UBD/Ub protein-protein interaction, including expanding on the phthalazinone scaffold and identification of quinazolinone analogues with improved predicted physicochemical properties and lead likeness.

## Acknowledgements

We thank past and present members of the Burslem lab for useful discussions and various contributions at the early stages of this project. We thank the University of Pennsylvania Post-Baccalaureate Research Education Program (PennPREP) for support. This work was funded by NIH NIGMS R35GM142505 to G.M.B.

## Table of Contents

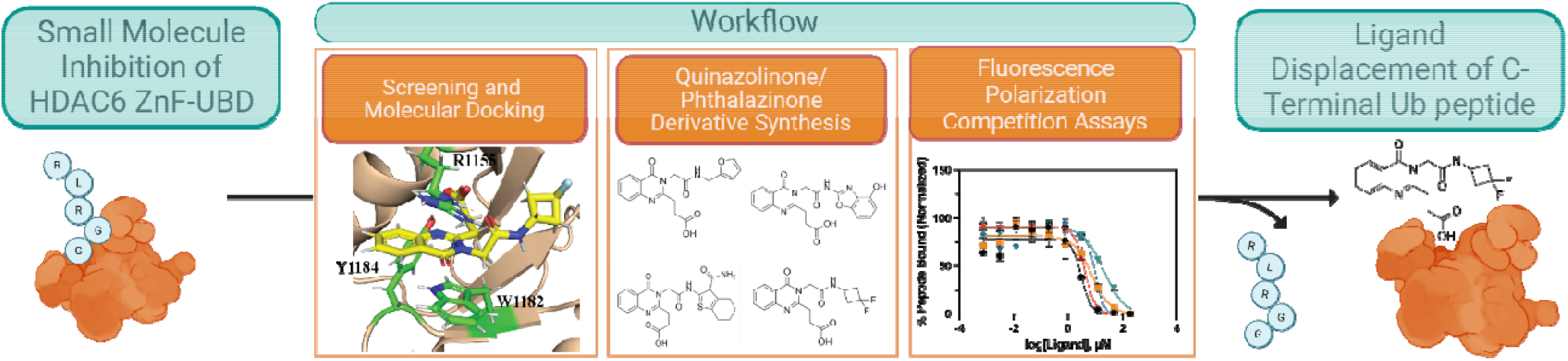

Here, we report the validation and expansion of small molecule inhibitors targeting the HDAC6 ZnF-UBD pocket. Following molecular modeling, hits were synthesized and candidate compounds were tested in fluorescence polarization of ubiquitin *C*-terminal peptide displacement. We additionally tested affinity probability scoring of the Boltz-2 co-folding system in comparison to our experimental results.

