## Supplementary Information for "Quinazolinone and Phthalazinone Inhibitors of the HDAC6/Ubiquitin Protein-Protein Interaction"

### Methods

### Chemical Synthesis:

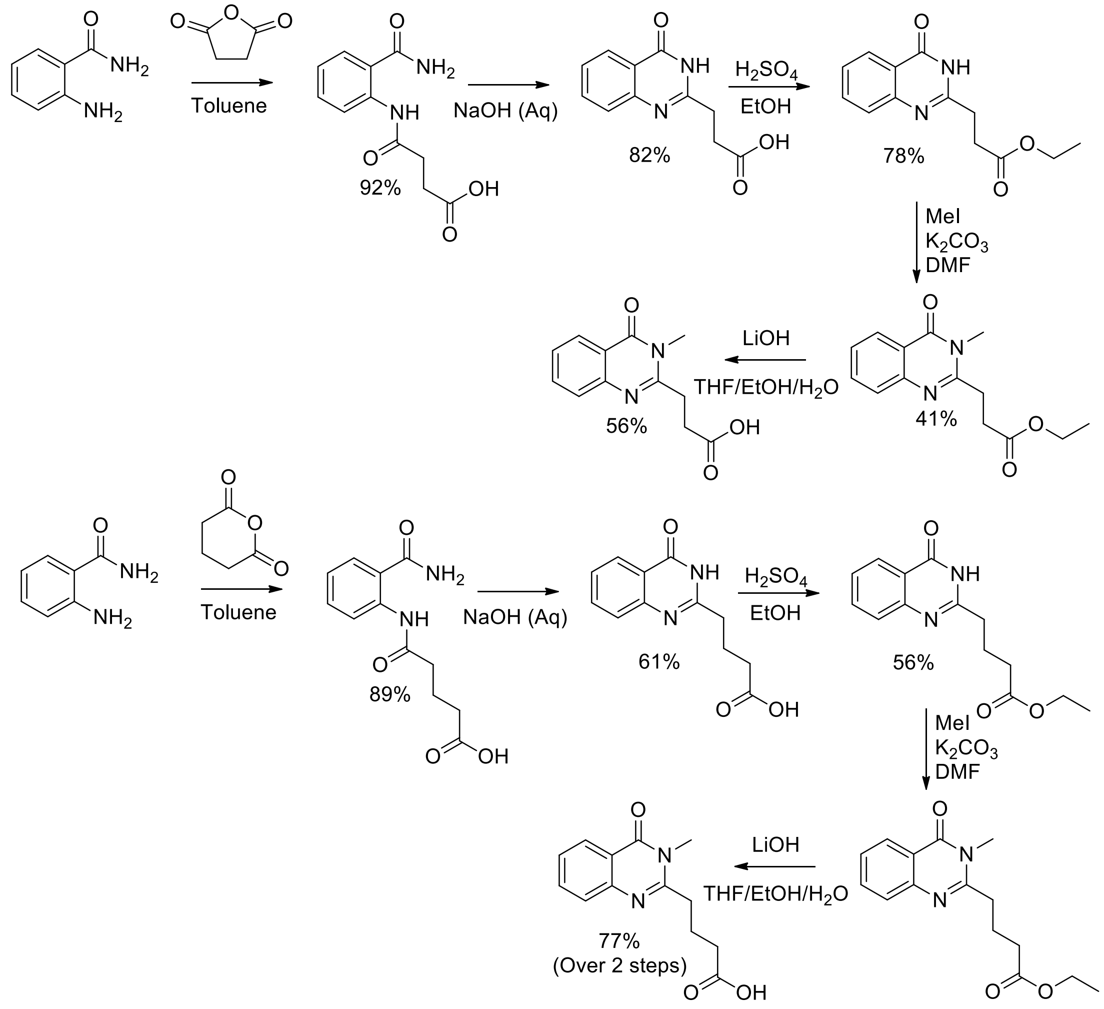


**Scheme 1 – Synthesis of Quinazolinone Cores**

**<u>4-((2-carbamoylphenyl)amino)-4-oxobutanoic acid:</u>**

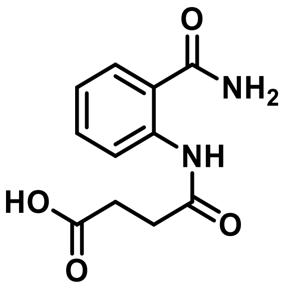

Anthranilamide (2.0 g, 14.7 mmol, 1 eq.) and toluene (25 mL) were added to a 100 mL round bottom flask equipped with a reflux condenser and stir-bar. Succinic anhydride (1.47 g, 14.7 mmol,

1 eq) was then added, and the reaction was heated to reflux for 3 hours. After cooling to room temperature, the product was collected via vacuum filtration, washed using cold diethyl ether (3 x 20 mL) and concentrated *in vacuo*, producing a white solid (3.2 g, 92%). *m/z* $[M + H]^+$ calcd for $C_{11}H_{12}N_2O_4$ 219.1; found: 219.1. Fragmented mass result comes from loss of water in LCMS positive mode. $^1$H NMR (400 MHz, DMSO) δ 12.16 (s, 1H), 11.73 (s, 1H), 8.44 (d, *J* = 8.3 Hz, 1H), 8.26 (s, 1H), 7.79 (d, *J* = 7.9 Hz, 1H), 7.72 (s, 1H), 7.47 (t, *J* = 7.8 Hz, 1H), 7.10 (t, *J* = 7.6 Hz, 1H), 2.55 (dd, *J* = 9.0, 4.6 Hz, 4H). $^{13}$C NMR (101 MHz, DMSO) δ 174.1, 171.2, 170.4, 141.5, 140.2, 132.7, 129.0, 122.7, 120.5, 32.5, 29.2.

**3-(4-oxo-3,4-dihydroquinazolin-2-yl) propanoic acid (Compound 1):**

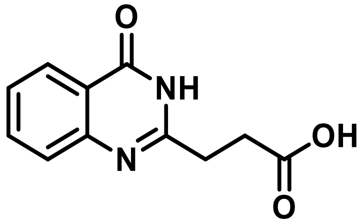


4-((2-carbamoylphenyl)amino)-4-oxobutanoic acid (3.2 g, 12 mmol, 1.0 eq) was mixed with a solution of sodium hydroxide (25 mL, 2.0 M) and heated to reflux for two hours. After cooling to room temperature, the resulting mixture was titrated using hydrochloric acid (1.0 M) until a pH between 5 and 6 was achieved. The resulting precipitate was collected via vacuum filtration and washed with cold water (3 x 20 mL). The product was then concentrated *in vacuo*, forming a white solid (2.5 g, 82%). *m/z* $[M + H]^+$ calcd for $C_{11}H_{10}N_2O_3$ 219.1; found: 219.1 $^1$H NMR (400 MHz, DMSO) δ 8.08 (dd, *J* = 8.0, 1.5 Hz, 1H), 7.78 (ddd, *J* = 8.5, 7.1, 1.6 Hz, 1H), 7.58 (dd, *J* = 8.3, 1.2 Hz, 1H), 7.47 (ddd, *J* = 8.1, 7.1, 1.2 Hz, 1H), 2.90 – 2.82 (m, 2H), 2.75 (td, *J* = 7.0, 1.2 Hz, 2H). $^{13}$C NMR (101 MHz, DMSO) 173.9, 162.0, 157.4, 148.0, 134.9, 126.7, 126.6, 126.3, 121.2, 30.4, 29.4

**Ethyl 3-(4-oxo-3,4-dihydroquinazolin-2-yl)propanoate:**

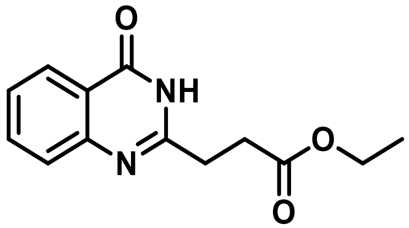


3-(4-oxo-3,4-dihydroquinazolin-2-yl) propanoic acid (2.5 g, 11.5 mmol, 1 eq) was suspended in ethanol (35 mL) and treated with sulfuric acid (2 mL). The reaction mixture was heated to reflux and stirred overnight until a turbid solution was observed. Additional monitoring by thin layer chromatography (1:1, Hex:EtOAc) was performed to ensure complete consumption of starting material. The reaction mixture was then concentrated, diluted with water (15 mL), and extracted using ethyl acetate (3 x 35 mL). The combined organic layers were dried over magnesium sulfate, filtered, and then concentrated *in vacuo* to produce a white solid (2.1 g, 78%). *m/z* $[M + H]^+$ calcd for $C_{13}H_{14}N_2O_3$ 247.1; found 247.1 $^1$H NMR (400 MHz, $CDCl_3$) δ 10.09 (s, 1H), 8.26 (dd, *J* = 8.0, 1.6 Hz, 1H), 7.74 (ddd, *J* = 8.5, 7.0, 1.5 Hz, 1H), 7.65 (d, *J* = 8.1 Hz, 1H), 7.50 – 7.42 (m, 1H), 4.20 (q, *J* = 7.1 Hz, 2H), 3.02 (dd, *J* = 7.2, 5.5 Hz, 2H), 2.90 (dd, *J* = 7.3, 5.5 Hz, 2H), 1.28 (t, *J* = 7.1 Hz, 3H).

**Ethyl 3-(3-methyl-4-oxo-3,4-dihydroquinazolin-2-yl)propanoate**:

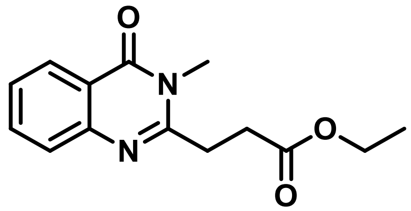


Ethyl 3-(4-oxo-3,4-dihydroquinazolin-2-yl)propanoate (0.2 g, 0.81 mmol, 1 eq), potassium carbonate (0.1 g, 0.97 mmol, 1.2 eq), and anhydrous DMF (15 mL) were added to a 50 mL round bottom flask equipped with a stir-bar. Methyl iodide (0.1 mL, 1.6 mmol, 2 eq) was then added dropwise and the mixture was stirred overnight. After consumption of starting material was confirmed by TLC (1:1, Hex:EtOAc) the reaction was diluted with water (15 mL) and extracted using ethyl acetate (3 x 15 mL). The combined organic layers were then washed with an aqueous solution of lithium chloride (5% w/v, 25 mL) and then brine (25 mL). The organic layers were dried over magnesium sulfate, filtered, and the crude product was purified by flash chromatography (10% to 35% Hex:EtOAc). The pure fractions were combined and concentrated *in vacuo*, producing a white solid (0.09 g, 41%). *m/z* $[M + H]^+$ calcd for $C_{14}H_{16}N_2O_3$ 261.1; found 261. $^1$H NMR (400 MHz, $CDCl_3$) δ 8.25 (dd, *J* = 8.1, 1.5 Hz, 1H), 7.74 – 7.66 (m, 1H), 7.58 (d, *J* = 8.1 Hz, 1H), 7.43 (t, *J* = 7.7 Hz, 1H), 4.19 (q, *J* = 7.1 Hz, 2H), 3.65 (s, 3H), 3.12 (t, *J* = 6.7 Hz, 2H), 2.95 (t, *J* = 6.7 Hz, 2H), 1.29 (t, *J* = 7.1 Hz, 3H).

**3-(3-methyl-4-oxo-3,4-dihydroquinazolin-2-yl)propanoic acid (Compound 31):**

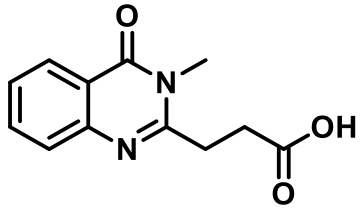


A solution of THF/EtOH/$H_2O$ (10 mL, 3:1:1) and Ethyl 3-(3-methyl-4-oxo-3,4-dihydroquinazolin-2-yl)propanoate (0.06 g, 0.23 mmol, 1 eq) were combined in a 25 mL round bottom flask equipped with a stir-bar. Lithium hydroxide (0.02 g, 0.92 mmol, 4 eq) was added and the reaction stirred overnight. After consumption of starting material was confirmed by TLC (1:1 Hex:EtOAc) the reaction was diluted with water (10 mL)and the organic solvents were removed via rotary evaporation. The remaining aqueous solution was titrated using hydrochloric acid (1M) until a pH between 6 and 7 was attained, forming an opaque white solid. The compound was then collected via vacuum filtration, washed with water and used without any additional purification. (0.13 g, 56%) *m/z* [M + H]$^+$ calcd for $C_{12}H_{12}N_2O_3$ 233.1; found 233.1. $^1$H NMR (400 MHz, DMSO) δ 8.09 (dd, *J* = 8.0, 1.6 Hz, 1H), 7.82 – 7.72 (m, 1H), 7.55 (d, *J* = 8.1 Hz, 1H), 7.46 (t, *J* = 7.5 Hz, 1H), 3.56 (s, 3H), 3.07 (t, *J* = 6.8 Hz, 2H), 2.74 (t, *J* = 6.8 Hz, 2H). $^{13}$C NMR (101 MHz, DMSO) δ 174.5, 161.8, 157.3, 147.2, 134.6, 127.2, 120.1, 30.9, 30.1, 29.8,

**5-((2-carbamoylphenyl)amino)-5-oxopentanoic acid:**

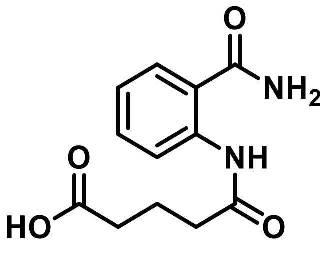


Anthranilamide (0.5 g, 3.67 mmol, 1 eq) and toluene (10 mL, 0.6 M) were added to a 50 mL round bottom flask equipped with a reflux condenser and stir-bar. Glutaric anhydride (0.42 g, 3.67 mmol, 1 eq) was then added, and the reaction was heated to reflux for 3 hours. After cooling to room temperature, the product was collected via vacuum filtration, washed using cold diethyl ether (3 x 10 mL) and concentrated *in vacuo*, producing a white solid (0.7 g, 76%). *m/z* [M + H]$^+$ calcd for $C_{12}H_{14}N_2O_4$ 233.1; found 233.4. Fragmented mass result comes from loss of water in LCMS positive mode. $^1$H NMR (400 MHz, DMSO) δ 12.08 (s, 1H), 11.66 (s, 1H), 8.45 (d, *J* = 8.3 Hz, 1H), 8.24 (s, 1H), 7.78 (dd, *J* = 7.9, 1.6 Hz, 1H), 7.69 (s, 1H), 7.50 – 7.06 (m, 2H), 2.38 (t, *J* = 7.5 Hz, 2H), 2.29 (t, *J* = 7.4 Hz, 2H), 1.82 (p, *J* = 7.5 Hz, 2H).

**4-(4-oxo-3,4-dihydroquinazolin-2-yl)butanoic acid:**

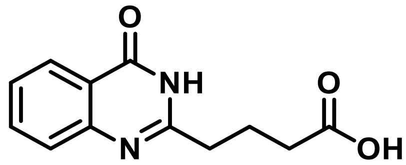


5-((2-carbamoylphenyl)amino)-5-oxopentanoic acid (0.5 g, 2 mmol, 1 eq) was mixed with a solution of sodium hydroxide (10 mL, 2.0 M) and heated to reflux for 2 hours. After cooling to room temperature, the resulting mixture was titrated using hydrochloric acid (1.0 M) until a pH between 5 and 6 was achieved. The resulting precipitate was collected via vacuum filtration and washed with cold water (3 x 20 mL). The resulting product formed was a white solid that was used without further purification (0.4 g, 81%). *m/z* [M + H]$^+$ calcd for $C_{12}H_{12}N_2O_3$ 215.2; found 215.6. Fragmented mass result comes from loss of water in LCMS positive mode. $^1$H NMR (400 MHz, DMSO) δ 8.07 (dd, *J* = 8.0, 1.5 Hz, 1H), 7.77 (ddd, *J* = 8.5, 7.1, 1.6 Hz, 1H), 7.60 (d, *J* = 8.1 Hz, 1H), 7.49 – 7.42 (m, 1H), 2.63 (t, *J* = 7.5 Hz, 2H), 2.31 (t, *J* = 7.4 Hz, 2H), 1.96 (p, *J* = 7.5 Hz, 2H).$^{13}$C NMR (101 MHz, $CDCl_3$) δ 173.09, 163.9, 155.6, 149.3, 134.8, 127.3, 126.3, 120.7, 34.8, 33.4, 29.7, 22.4.

**Ethyl 4-(4-oxo-3,4-dihydroquinazolin-2-yl)butanoate:**

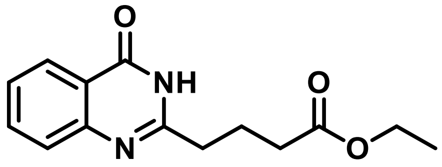

4-(4-oxo-3,4-dihydroquinazolin-2-yl)butanoic acid (0.2 g, 0.86 mmol, 1eq) was suspended in ethanol (15 mL) and treated with sulfuric acid (3 mL). The reaction mixture was heated to reflux and stirred overnight until a turbid solution was observed. Additional monitoring by thin layer chromatography (1:1, Hex:EtOAc) was performed to ensure complete consumption of starting material. The reaction mixture was then concentrated, diluted with water (25 mL), and extracted using ethyl acetate (3 x 35 mL). The combined organic layers were dried over magnesium sulfate, filtered, and then concentrated *in vacuo* to produce a white solid (0.13g, 56%). *m/z* $[M + H]^+$ calcd for $C_{14}H_{16}N_2O_3$ 261.1; found 261.1$^1$H NMR (400 MHz, $CDCl_3$) δ 11.49 (s, 1H), 8.29 (dd, *J* = 8.0, 1.5 Hz, 1H), 7.76 (ddd, *J* = 8.5, 7.0, 1.6 Hz, 1H), 7.69 (dd, *J* = 8.1, 1.2 Hz, 1H), 7.47 (ddd, *J* = 8.2, 7.0, 1.3 Hz, 1H), 4.13 (q, *J* = 7.1 Hz, 2H), 2.84 (t, *J* = 7.5 Hz, 2H), 2.50 (t, *J* = 7.3 Hz, 2H), 2.22 (p, *J* = 7.3 Hz, 2H), 1.25 (t, *J* = 7.1 Hz, 3H).$^{13}$C NMR (101 MHz, $CDCl_3$) δ 173.1, 163.9, 149.3, 134.8, 127.3, 126.5, 126.3, 120.7, 60.6, 34.78, 33.3, 29.7, 22.4, 14.2.

**4-(3-methyl-4-oxo-3,4-dihydroquinazolin-2-yl)butanoic acid (Compound 2):**

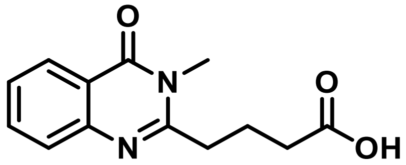

Ethyl 4-(4-oxo-3,4-dihydroquinazolin-2-yl)butanoate (0.05 g, 0.153 mmol, 1 eq), potassium carbonate (0.03 g, 0.184 mmol, 1.2 eq), and anhydrous DMF (5 mL) were added to a 25 mL round bottom flask equipped with a stir-bar. Methyl iodide (0.02 mL, 0.31 mmol, 2 eq) was then added dropwise and the mixture was stirred overnight. After consumption of starting material was confirmed by TLC (1:1, Hex:EtOAc) the reaction was diluted with water and extracted using ethyl acetate (3 x 15 mL). The combined organic layers were then washed with an aqueous solution of lithium chloride (5% w/v, 25 mL) and then brine (25 mL). The organic layers were dried over magnesium sulfate, filtered, and the crude product was taken to the next step. THF/MeOH (8 mL, 1:1) and the crude product were combined in a 25 mL round bottom flask equipped with a stir-bar. Lithium hydroxide (1 mL, 1 M, 0.84 mmol, 1.5 eq) was added and the reaction stirred overnight. After consumption of starting material was confirmed by TLC (1:1 Hex:EtOAc) the reaction was diluted with water and the organic solvents were evaporated. The remaining aqueous solution was titrated using hydrochloric acid (1M) until a pH between 6 and 7 was attained, forming an opaque white solid. The resulting precipitate was collected via vacuum filtration and washed with cold water (3 x 20 mL). The product was then concentrated *in vacuo*, forming a white solid (0.014 g,

77%). HRMS $[M + H]^+$ calcd for $C_{13}H_{14}N_2O_3$ 247.1082 found 247.1072; Δppm = 2.08. $^1$H NMR (400 MHz, DMSO) δ 12.08 (s, 1H), 8.10 (dd, $J$ = 8.0, 1.5 Hz, 1H), 7.77 (ddd, $J$ = 8.5, 7.1, 1.6 Hz, 1H), 7.60 (dd, $J$ = 8.2, 1.1 Hz, 1H), 7.47 (ddd, $J$ = 8.1, 7.0, 1.2 Hz, 1H), 3.53 (s, 3H), 2.88 (dd, $J$ = 8.1, 7.0 Hz, 2H), 2.41 (t, $J$ = 7.2 Hz, 2H), 2.00 (p, $J$ = 7.3 Hz, 2H).

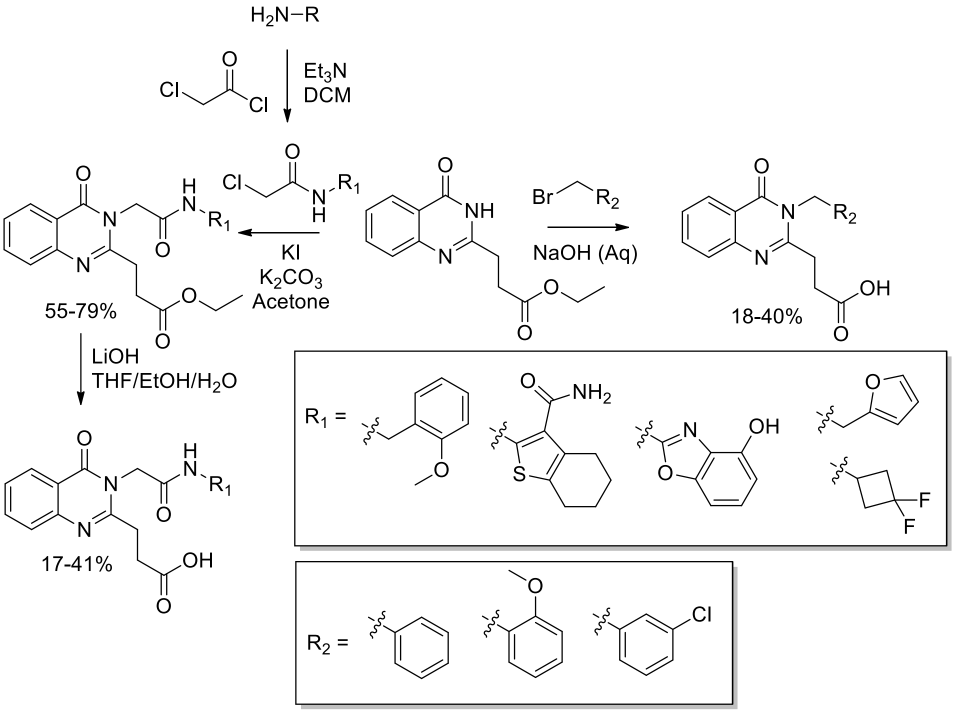


**Scheme 2 – Synthesis of Quinazolinone Derivatives**

**3-(3-benzyl-4-oxo-3,4-dihydroquinazolin-2-yl)propanoic acid (Compound 3):**

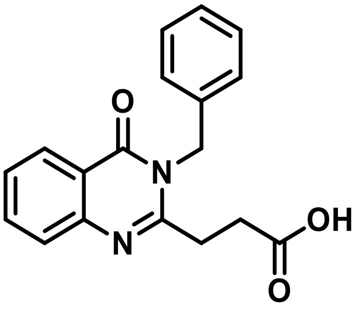

Ethyl 3-(4-oxo-3,4-dihydroquinazolin-2-yl)propanoate (0.025 g, 0.1 mmol, 1 eq), sodium hydroxide (0.02g, 0.5mmol, 5eq) and DMF (10 mL) were added to a 50 mL round bottom flask and stirred for 30 minutes. Benzyl bromide (0.2 g, 0.112 mmol, 1.1 eq) was then added dropwise and the mixture was heated to 50° C for 2 hours. After cooling to room temperature, the reaction was diluted with water (10 mL) and the pH was titrated to between 2-3 using acetic acid. The compound was extracted using ethyl acetate (3 x 25 mL), then brine. The organic layers were combined, dried over magnesium sulfate and concentrated *in vacuo*. The resulting compound was then recrystallized in methanol to produce a white solid (0.03 g, 78%). HRMS $[M + H]^+$ calcd for $C_{18}H_{16}N_2O_3$ 309.1161; found 309.1236; Δppm = 0.63. $^1H$ NMR (400 MHz, $CDCl_3$) δ 8.06 – 7.71 (m, 1H), 7.64 – 7.46 (m, 1H), 7.45 – 7.25 (m, 3H), 7.07 – 6.87 (m, 4H), 4.61 (s, 2H), 3.93 (s, 2H), 3.86 (t, $J$ = 4.5 Hz, 2H). $^{13}C$ NMR (101 MHz, $CDCl_3$) δ 175.7, 163.1, 162.4, 155.4, 146.8, 135.7, 134.4, 129.0, 128.4, 127.7, 127.0, 126.7, 126.5, 120.3, 36.7, 31.6, 20.8.

**3-(3-(3-chlorobenzyl)-4-oxo-3,4-dihydroquinazolin-2-yl)propanoic acid (Compound 4):**

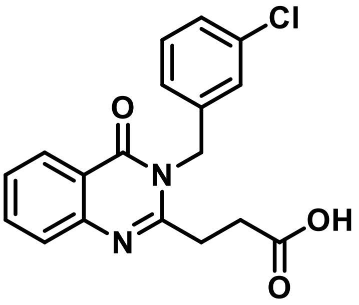


Ethyl 3-(4-oxo-3,4-dihydroquinazolin-2-yl)propanoate (0.075 g, 0.3 mmol, 1 eq.), sodium hydroxide (0.06 g, 1.5 mmol, 5 eq.) and DMF (10 mL) were added to a 50 mL round bottom flask and stirred for 30 minutes. 1-(bromomethyl)-3-chlorobenzene (0.07 g, 0.33 mmol, 1.1 eq.) was then added dropwise and the mixture was heated to 50° C for 2 hours. After cooling to room temperature, the reaction was diluted with water (10 mL) and the pH was titrated to between 2-3 using acetic acid. The compound was extracted using ethyl acetate (3 x 25 mL), then brine. The organic layers were combined, dried over magnesium sulfate and concentrated *in vacuo*. The resulting compound was then recrystallized in methanol to produce a white solid (0.014g, 14%). HRMS $[M + H]^+$ calcd for $C_{18}H_{15}ClN_2O_3$ 343.0849; found 343.0842; Δppm = -0.71. $^1H$ NMR (400 MHz, DMSO) δ 8.20 – 8.05 (m, 1H), 7.68 – 7.47 (m, 3H), 7.49 – 7.24 (m, 4H), 5.41 (s, 2H), 2.98 (t, 2H), 2.74 (t, 2H). $^{13}C$ NMR (101 MHz, DMSO) δ 174.1, 161.9, 156.7, 147.1, 139.5, 135.1, 131.1, 130.4, 127.8, 127.4, 127.1, 127.0, 126.9, 126.8, 126.5, 125.4, 30.2, 29.3.

**3-(3-(2-methoxybenzyl)-4-oxo-3,4-dihydroquinazolin-2-yl)propanoic acid (Compound 5):**

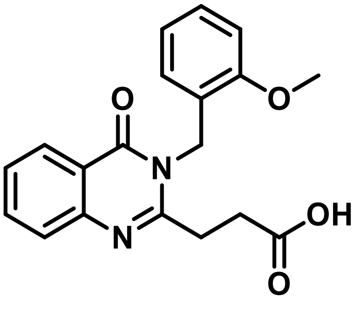


Ethyl 3-(4-oxo-3,4-dihydroquinazolin-2-yl)propanoate (0.1 g, 0.41 mmol, 1 eq.), sodium hydroxide (0.08 g, 2 mmol, 5 eq.) and DMF (10 mL) were added to a 50 mL round bottom flask and stirred for 30 minutes. 1-(bromomethyl)-3-chlorobenzene (0.085g, 0.45 mmol, 1.1 eq.) was then added dropwise and the mixture was heated to 50° C for 2 hours. After cooling to room temperature, the reaction was diluted with water (10 mL) and the pH was titrated to between 2-3 using acetic acid. The compound was extracted using ethyl acetate (3 x 25 mL), then brine. The organic layers were combined, dried over magnesium sulfate and concentrated *in vacuo*. The resulting compound was then recrystallized in methanol to produce a white solid (0.02 g, 15%). HRMS $[M + H]^+$ calcd for $C_{19}H_{18}N_2O_4$ 339.1344; found 339.1338; Δppm -0.50. $^1$H NMR (400 MHz, $CDCl_3$) δ 7.51 – 7.42 (m, 2H), 7.40 – 7.35 (m, 2H), 7.20 – 7.13 (m, 4H), 5.21 (s, 2H), 3.76 (s, 3H), 3.52 (t, 2H), 3.34 (t, 2H). $^{13}$C NMR (101 MHz, $CDCl_3$) δ 172.7, 170.1, 169.3, 160.8, 138.1, 136.5, 135.2, 135.1, 128.5, 128.2, 127.8, 126.9, 126.4, 124.1, 121.7, 114.4, 62.4, 52.3, 41.9.

**Ethyl 3-(3-(2-((2-methoxybenzyl)amino)-2-oxoethyl)-4-oxo-3,4-dihydro quinazolin-2-yl)propanoate (Compound 6):**

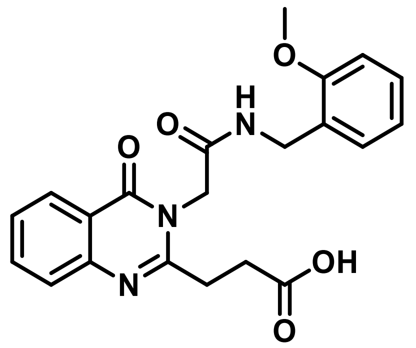


Potassium iodide (0.13 g, 0.93 mmol, 1.5 eq), 2-chloro-N-(2-methoxybenzyl)acetamide (0.13 g, 0.6 mmol, 1.5 eq), and acetone (4 mL) were combined in a round bottom flask equipped with a reflux condenser and stir bar. The reaction stirred for 30 minutes before potassium carbonate (0.09 g, 0.6 mmol, 1.5 eq) and Ethyl 3-(4-oxo-3,4-dihydroquinazolin-2-yl)propanoate in acetone (0.2 g, 0.82 mmol, 1 eq) were added. The mixture was then heated to reflux for 4 hours. After cooling to room temperature, the resulting solution was diluted with water (25 mL) and extracted using ethyl acetate (3 x 25 mL). The combined organic layers were then washed with water and brine before being concentrated. The crude product was then added to a solution of THF/MeOH (8 mL, 1:1), lithium hydroxide (1 mL, 1M), and stirred overnight. The reaction was then diluted with water and titrated using hydrochloric acid (1M) until a pH below 3 was attained, resulting in an opaque white solid. The compound was then collected vacuum filtration and washed with water without any additional purification. (0.02 g, 11%). HRMS $[M + Na]^+$ calcd for $C_{21}H_{21}N_3O_5$ 418.1379; found 418.1369; Δppm -1.14. $^1$H NMR (400 MHz, DMSO) δ 8.66 (t, $J$ = 5.8 Hz, 1H), 8.10 (dd, $J$ = 8.0, 1.5 Hz, 1H), 7.81 (ddd, $J$ = 8.6, 7.1, 1.5 Hz, 1H), 7.59 (d, $J$ = 8.1 Hz, 1H), 7.50 (t, $J$ = 7.6 Hz, 1H), 7.25 (dd, $J$ = 9.4, 6.9 Hz, 1H), 7.04 – 6.90 (m, 2H), 4.89 (s, 2H), 4.29 (d, $J$ = 5.6 Hz, 2H), 3.80 (s, 3H), 3.01 (t, $J$ = 6.8 Hz, 2H), 2.76 (t, $J$ = 6.8 Hz, 2H).$^{13}$C NMR (101 MHz, DMSO) δ 174.1, 167.1, 161.7, 157.2, 156.8, 147.2, 134.9, 128.7, 128.4, 127.3, 127.0, 126.8, 126.7, 120.7, 120.2, 111.0, 55.8, 45.7, 38.0, 30.5, 29.3.

**3-(3-(2-((furan-2-ylmethyl)amino)-2-oxoethyl)-4-oxo-3,4-dihydroquinazolin-2-yl)propanoic acid (Compound 9):**

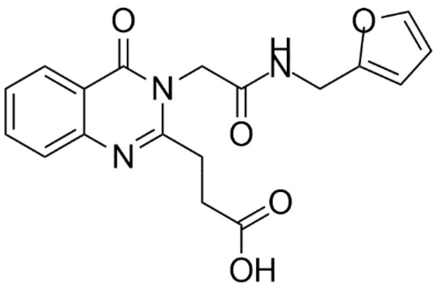

Potassium iodide (0.15 g, 0.91 mmol, 1.5 eq), 2-chloro-N-(furan-2-ylmethyl)acetamide (0.16 g, 0.91 mmol, 1.5 eq), and acetone (4 mL) were combined in a round bottom flask equipped with a reflux condenser and stir-bar. The reaction stirred for 30 minutes before potassium carbonate (0.13 g, 0.91 mmol, 1.5 eq) and Ethyl 3-(4-oxo-3,4-dihydroquinazolin-2-yl)propanoate (0.15 g, 0.61 mmol, 1 eq) in acetone (6 mL) were added. The mixture was then heated to reflux for 4 hours. After cooling to room temperature, the solution was diluted with water (25 mL) and extracted using ethyl acetate (3 x 25 mL). The product was then purified via flash chromatography (20% to 45% Hex:EtOAc) to produce a brown oil (0.16 g, 51%). The product was then added to a solution of THF/MeOH (8 mL, 1:1), lithium hydroxide (1 mL, 1M), and stirred overnight. The reaction was then diluted with water and titrated using hydrochloric acid (1M) until a pH below 3 was attained, resulting in an opaque white solid. The compound was then collected vacuum filtration and washed with water and further purified using semi-preparative HPLC. HRMS $[M + Na]^+$ calcd for $C_{18}H_{17}N_3O_5$ 378.106592; found 378.1041; Δppm -5.25. $^1H$ NMR (400 MHz, $CDCl_3$) δ 8.27 (dd, *J* = 8.0, 1.8 Hz, 1H), 8.21 (dd, *J* = 8.0, 1.8 Hz, 1H), 7.75 (tdd, *J* = 8.1, 7.0, 1.6 Hz, 2H), 7.67 (dd, *J* = 8.3, 1.4 Hz, 1H), 7.61 (dd, *J* = 8.3, 1.4 Hz, 1H), 7.46 (qd, *J* = 8.0, 1.3 Hz, 2H), 4.82 (s, 2H), 4.21 – 4.17 (m, 3H), 4.13 (q, *J* = 7.2 Hz, 5H), 3.02 – 2.97 (m, 2H).13C NMR (101 MHz, $CDCl_3$) δ 173.10, 172.89, 171.18, 166.96, 163.19, 162.67, 154.76, 154.54, 149.09, 147.03, 127.26, 126.84, 126.70, 126.57, 126.35, 120.84, 119.90, 60.40, 31.59, 22.63 (d, *J* = 4.4 Hz), 21.04, 14.23 (d, *J* = 7.6 Hz), 11.42.

**3-(3-(2-((3,3-difluorocyclobutyl)amino)-2-oxoethyl)-4-oxo-3,4-dihydroquinazolin-2-yl)propanoic acid (Compound 10):**

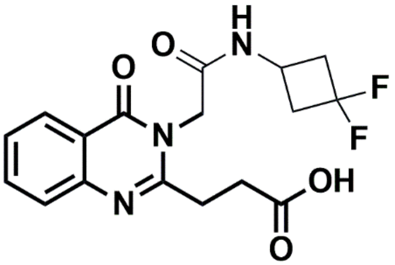

Potassium iodide (45 mg, 0.27 mmol, 1.5 eq), 2-chloro-N-(3,3-difluorocyclobutyl)acetamide (132 mg, 0.72 mmol, 4 eq), and acetone (4 mL) were combined in a round bottom flask equipped with a reflux condenser and stir-bar. The reaction stirred for 30 minutes before potassium carbonate (37

mg, 0.27 mmol, 1.5 eq) and Ethyl 3-(4-oxo-3,4-dihydroquinazolin-2-yl)propanoate (25 mg, 0.18 mmol, 1 eq) in acetone (6 mL) were added. The mixture was then heated to reflux for 4 hours. After cooling to room temperature, the solution was diluted with water (25 mL) and extracted using ethyl acetate (3 x 25 mL). The crude product was then added to a solution of THF/MeOH (8 mL, 1:1), lithium hydroxide (1 mL, 1M), and stirred overnight. The reaction was then diluted with water and titrated using hydrochloric acid (1M) until a pH below 3 was attained, resulting in an opaque white solid. The compound was then collected vacuum filtration and washed with water and further purified using semi-preparative HPLC to afford Compound 10 as an opaque white solid (18 mg, 32%). HRMS $[M + Na]^+$ calcd for $C_{17}H_{17}F_2N_3O_4$ 388.1084; found 388.1075; Δppm -1.00. $^1$H NMR (400 MHz, $CDCl_3$) δ 7.28 (m, 2H), 7.00 (m, 2H), 3.77 (s, 3H), 1.88 (p, $J$ = 3.1 Hz, 5H), 1.45 (s, 3H), 1.31 – 1.23 (m, 3H), 0.94 – 0.82 (m, 1H). $^{13}$C NMR (101 MHz, $CDCl_3$) δ 166.73, 162.41, 150.97, 143.22, 142.24, 110.48, 107.42, 67.97, 53.43, 36.34, 34.55, 30.94, 30.33, 29.71, 25.60. $^{19}$F NMR (376 MHz, $CDCl_3$) δ -84.09 – -85.26 (m), -97.79 (dd, $J$ = 226.9, 199.0 Hz).

**3-(3-(2-((3-carbamoyl-4,5,6,7-tetrahydrobenzo[b]thiophen-2-yl)amino)-2-oxoethyl)-4-oxo-3,4-dihydroquinazolin-2-yl)propanoic acid (Compound 7):**

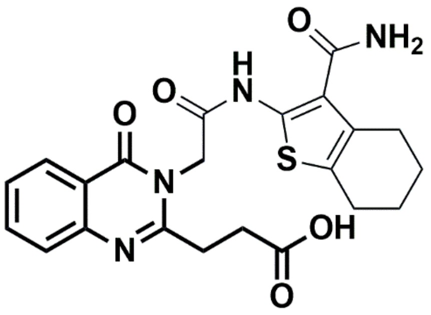

Potassium iodide (23 mg, 0.14 mmol, 1.5 eq), 2-(2-chloroacetamido)-4,5,6,7-tetrahydrobenzo[b]thiophene-3-carboxamide (73 mg, 0.27 mmol, 3 eq), and acetone (4 mL) were combined in a round bottom flask equipped with a reflux condenser and stir-bar. The reaction stirred for 30 minutes before potassium carbonate (19 mg, 0.14 mmol, 1.5 eq) and Ethyl 3-(4-oxo-3,4-dihydroquinazolin-2-yl)propanoate (23 mg, 0.09 mmol, 1 eq) in acetone (6 mL) were added. The mixture was then heated to reflux for 4 hours. After cooling to room temperature, the solution was diluted with water (25 mL) and extracted using ethyl acetate (3 x 25 mL). The crude product was then added to a solution of THF/MeOH (8 mL, 1:1), lithium hydroxide (1 mL, 1M), and stirred overnight. The reaction was then diluted with water and titrated using hydrochloric acid (1M) until a pH below 3 was attained, resulting in an opaque white solid. The compound was then collected vacuum filtration and washed with water and further purified using semi-preparative HPLC to afford Compound 7 as a dark orange solid (18 mg, 41%). HRMS $[M + H]^+$ calcd for $C_{22}H_{22}N_4O_5S$ 455.1389; found 455.1381; Δppm -0.60. $^1$H NMR (400 MHz, CDCl3) δ 8.29 – 8.24 (m, 1H), 7.76 (ddd, $J$ = 8.6, 7.1, 1.6 Hz, 1H), 7.67 (dt, $J$ = 8.3, 1.1 Hz, 2H), 7.47 (ddd, $J$ = 8.1, 7.0, 1.3 Hz, 1H), 4.20 (q, $J$ = 7.1 Hz, 2H), 3.14 – 3.07 (m, 2H), 2.98 – 2.93 (m, 2H), 2.19 (s, 2H), 1.28 (t, $J$ = 7.1

Hz, 4H). $^{13}C$ NMR (101 MHz, CDCl3) δ 172.96, 163.51, 154.88, 149.13, 134.63, 127.24, 126.33, 120.89, 61.02, 53.44, 30.94, 30.87, 30.17, 26.46, 24.38, 22.83, 22.59, 14.20.

**3-(3-(2-((4-hydroxybenzo[d]oxazol-2-yl)amino)-2-oxoethyl)-4-oxo-3,4-dihydroquinazolin-2-yl)propanoic acid (Compound 8):**

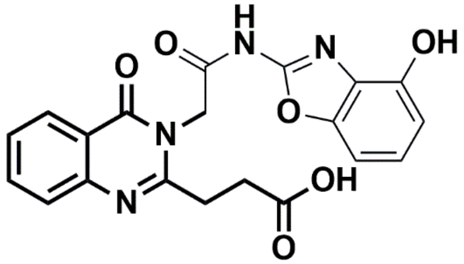


Potassium iodide (14 mg, 0.09 mmol, 1.5 eq), 2-chloro-N-(4-hydroxybenzo[d]oxazol-2-yl)acetamide (90 mg, 0.39 mmol, 6.5 eq), and acetone (4 mL) were combined in a round bottom flask equipped with a reflux condenser and stir-bar. The reaction stirred for 30 minutes before potassium carbonate (12 mg, 0.09 mmol, 1.5 eq) and Ethyl 3-(4-oxo-3,4-dihydroquinazolin-2-yl)propanoate (15 mg, 0.06 mmol, 1 eq) in acetone (6 mL) were added. The mixture was then heated to reflux for 4 hours. After cooling to room temperature, the solution was diluted with water (25 mL) and extracted using ethyl acetate (3 x 25 mL). The crude product was then added to a solution of THF/MeOH (8 mL, 1:1), lithium hydroxide (1 mL, 1M), and stirred overnight. The reaction was then diluted with water and titrated using hydrochloric acid (1M) until a pH below 3 was attained, resulting in an opaque white solid. The compound was then collected vacuum filtration and washed with water and further purified using semi-preparative HPLC to afford (17 mg, 67%). HRMS $[M + Na]^+$ calcd for $C_{20}H_{16}N_4O_6$ 431.0968; found 431.0972; Δppm 2.42. $^{1}H$ NMR (400 MHz, $CDCl_3$) δ 8.32 – 8.24 (m, 1H), 7.76 (ddd, $J$ = 8.6, 7.1, 1.6 Hz, 1H), 7.67 (dt, $J$ = 8.3, 1.1 Hz, 2H), 7.47 (ddd, $J$ = 8.1, 7.0, 1.3 Hz, 1H), 4.20 (q, $J$ = 7.1 Hz, 2H), 3.14 – 3.09 (m, 1H), 3.02 – 2.87 (m, 2H), 2.79 – 2.62 (m, 1H), 1.96 – 1.77 (m, 1H), 1.28 (t, $J$ = 7.1 Hz, 3H). $^{13}C$ NMR (101 MHz, $CDCl_3$) δ 172.96, 163.38, 154.88, 149.13, 134.63, 127.24, 126.42 (d, $J$ = 17.4 Hz), 120.89, 61.02, 53.44, 30.87, 30.17, 26.46, 24.38, 22.71 (d, $J$ = 25.1 Hz), 14.20, 7.97.

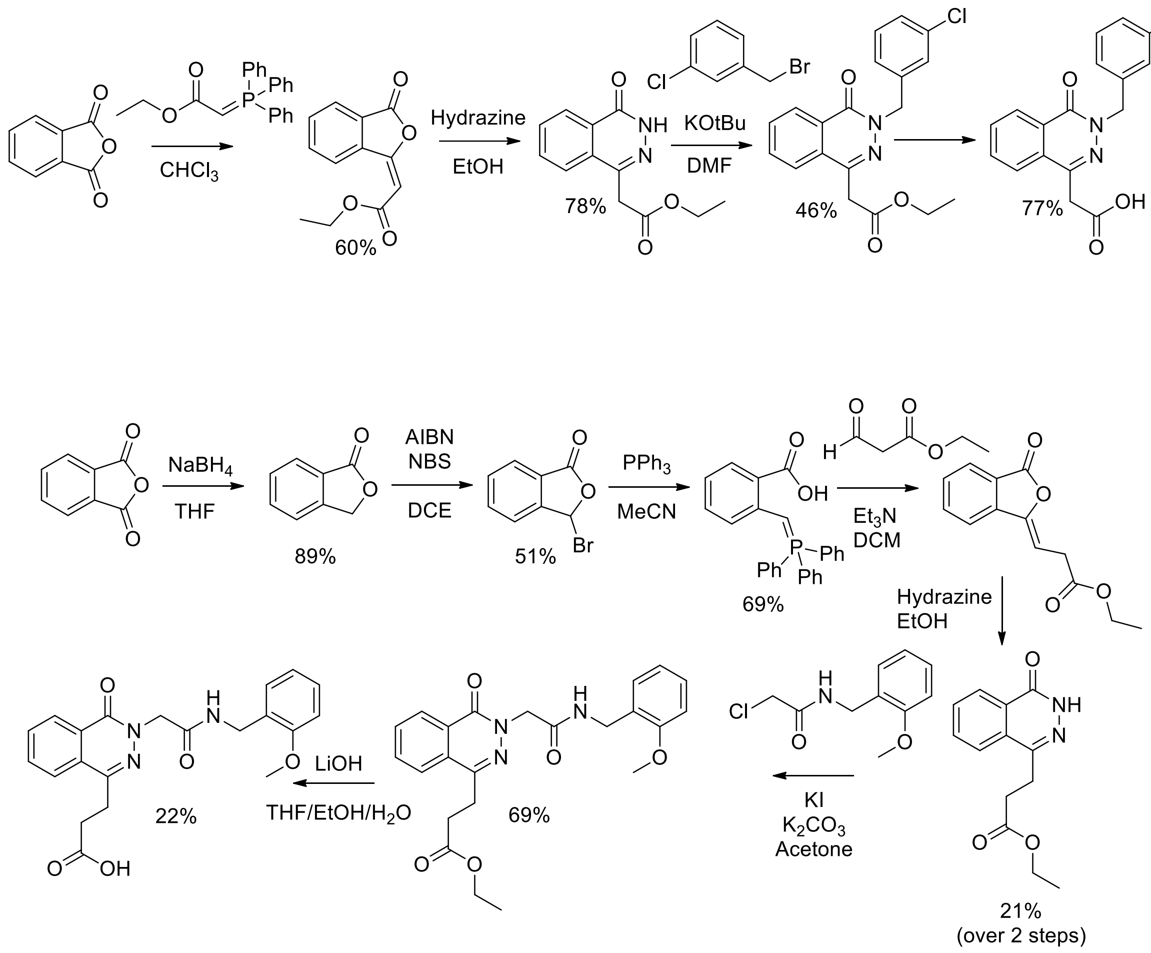


**Scheme 3 - Synthesis of Phthalazinones**

**<u>Ethyl (E)-2-(3-oxoisobenzofuran-1(3H)-ylidene)acetate:</u>**

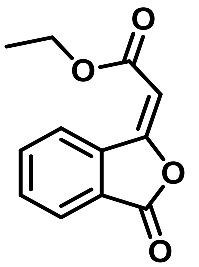

Chloroform (10 mL) and phthalic anhydride (0.4 g, 2.87 mmol, 1 eq) were combined in a 50 mL round bottom flask equipped with a reflux condenser and stir-bar. (Carbethoxymethylene) triphenylphosphorane (1g, 2.87 mmol, 1 eq) was then added portion-wise to the flask and the reaction was heated to reflux for 3 hours. Consumption of starting material was monitored by TLC (3:1 Hex:EtOAc) before the mixture was concentrated *in vacuo* and then purified by flash chromatography (5% to 15% Hex:EtOAc, 0.4 g, 63%). *m/z* $[M + H]^+$ calcd for $C_{12}H_{10}O_4$ 219.0;

found 219.0 $^{1}H$ NMR (400 MHz, $CDCl_3$) δ 9.06 (dd, $J$ = 8.0, 1.1 Hz, 1H), 7.98 (dd, $J$ = 7.6, 1.1 Hz, 1H), 7.87 – 7.80 (m, 1H), 7.72 (td, $J$ = 7.5, 1.0 Hz, 1H), 6.16 (d, $J$ = 0.9 Hz, 1H), 4.31 (qd, $J$ = 7.2, 0.9 Hz, 2H), 1.38 (td, $J$ = 7.2, 0.9 Hz, 3H). $^{13}C$ NMR (101 MHz, $CDCl_3$) δ 165.7, 165.6, 157.8, 136.2, 135.3, 132.5, 128.2, 126.6, 125.4, 102.5, 60.9, 14.3.

**Ethyl 2-(4-oxo-3,4-dihydrophthalazin-1-yl)acetate:**

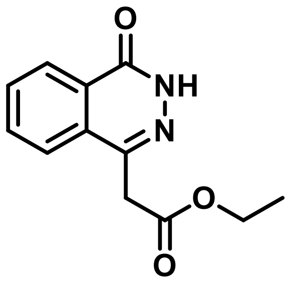


Ethyl (E)-2-(3-oxoisobenzofuran-1(3H)-ylidene)acetate (0.3 g, 0.125 mmol, 1 eq) was dissolved in ethanol (10 mL) in a 25 mL round bottom flask equipped with a stir-bar. Hydrazine hydrate (0.22 mL, 0.125 mmol, 1 eq) was added dropwise and the reaction was heated to 50° C for 2 hours. The reaction was then cooled to room temperature where the product was collected via vacuum filtration and washed with ethanol (0.28 g, 96%). $m/z$ $[M + H]^+$ calcd for $C_{12}H_{12}N_2O_3$ 233.1; found 233.1. $^{1}H$ NMR (400 MHz, DMSO) δ 12.65 (d, $J$ = 2.8 Hz, 1H), 8.26 (dd, $J$ = 7.7, 2.7 Hz, 1H), 7.93 (ddd, $J$ = 8.8, 5.2, 2.2 Hz, 1H), 7.85 (dt, $J$ = 7.8, 4.6 Hz, 2H), 4.11 (tt, $J$ = 9.1, 4.6 Hz, 2H), 4.05 (d, 2H), 1.16 (td, $J$ = 7.1, 2.4 Hz, 3H). $^{13}C$ NMR (101 MHz, DMSO) δ 170.3, 159.9, 141.4, 134.0, 132.1, 129.9, 128.0, 126.4, 125.9, 61.2, 38.4, 14.5.

**Ethyl 2-(3-(3-chlorobenzyl)-4-oxo-3,4-dihydrophthalazin-1-yl)acetate:**

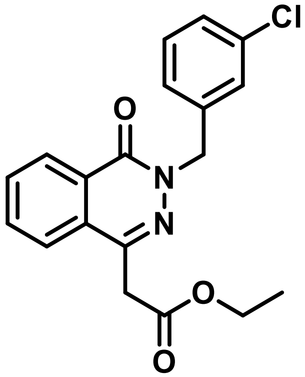


Ethyl 2-(4-oxo-3,4-dihydrophthalazin-1-yl)acetate (0.08 g, 0.32 mmol, 1 eq), DMF (5 mL), and potassium tert-butoxide (0.05 g, 0.36 mmol, 1.1 eq) were added to a 25 mL round bottom flask equipped with a stir-bar. The reaction was stirred for 30 minutes at room temperature, after which the colorless mixture became an opaque yellow color. 1-(bromomethyl)-2-chlorobenzene (0.07 g, 0.32 mmol, 1 eq) was then added dropwise to the mixture and the reaction stirred overnight. Upon confirming complete consumption of starting material via TLC (4:1 Hex:EtOAc), the reaction was diluted with ethyl acetate (30 mL), washed with a lithium chloride solution (5% w/v, 30 mL) and then brine. The organic layer was then concentrated *in vacuo* and the crude product underwent purification using flash chromatography (5% to 25% Hex:EtOAc) (0.06g, 46%). *m/z* $[M + H]^+$ calcd for $C_{19}H_{17}ClN_2O_3$ 357.1; found 357.1 $^1H$ NMR (400 MHz, $CDCl_3$) δ 8.47 (dd, *J* = 7.7, 1.7 Hz, 1H), 7.85 – 7.73 (m, 2H), 7.70 (dd, *J* = 7.5, 1.6 Hz, 1H), 7.42 (s, 1H), 7.33 (td, *J* = 5.5, 3.0 Hz, 1H), 7.25 – 7.21 (m, 2H), 5.35 (s, 2H), 4.19 (q, *J* = 7.1 Hz, 2H), 3.97 (s, 2H), 1.23 (t, *J* = 7.1 Hz, 3H).

**2-(3-(3-chlorobenzyl)-4-oxo-3,4-dihydrophthalazin-1-yl)acetic acid (Compound 11):**

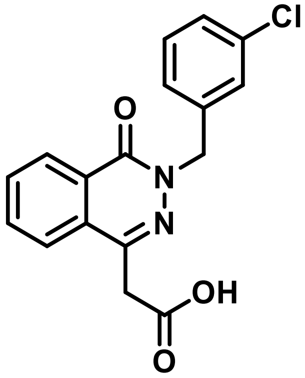


A solution of THF/MeOH (8 mL, 1:1) and Ethyl 2-(4-oxo-3,4-dihydrophthalazin-1-yl)acetate (0.03 g, 0.056 mmol, 1 eq) were combined in a 25 mL round bottom flask equipped with a stir-bar. Lithium hydroxide (0.02 g, 0.084 mmol, 1.5 eq) was added and the reaction stirred overnight. The reaction was diluted with water (10 mL) and the organic solvents were removed via rotary evaporation. The remaining aqueous solution was titrated using hydrochloric acid (1 M) until a pH below 3 was attained, forming an opaque white solid. The compound was then collected by vacuum filtration, washed with water and used without any additional purification (0.014 g, 77%). *m/z* $[M + H]^+$ calcd for $C_{17}H_{13}ClN_2O_3$ 329.1; found 329.1. $^1H$ NMR (400 MHz, $CDCl_3$) δ 8.47 (dd, *J* = 7.6, 1.7 Hz, 1H), 7.84 – 7.73 (m, 2H), 7.70 (dd, *J* = 7.5, 1.6 Hz, 1H), 7.42 (s, 1H), 7.36 – 7.31 (m, 1H), 7.24 – 7.22 (m, 2H), 5.35 (s, 2H), 3.97 (s, 2H).$^{13}C$ NMR (101 MHz, CDCl3) δ 169.7, 159.3, 141.0, 139.0, 134.4, 133.4, 131.8, 129.9, 129.5, 128.6, 128.2, 128.0, 127.6, 126.9, 124.7, 54.2, 39.0

**2-((triphenyl-l5-phosphaneylidene)methyl)benzoic acid:**

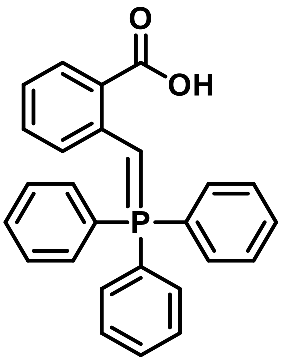

6-methoxy-*3H*-isobenzofuran-1-one (prepared as described in *Chem. Asian J.* **2019**, *14*, 1278) (0.5g, 2.34 mmol, 1 eq.) was suspended in acetonitrile (10 mL) in a 50 mL round bottom flask equipped with a reflux condenser and stir-bar. Triphenylphosphine (0.92 g, 3.52mmol, 1.5 eq.) was added and the mixture was heated to reflux for 3 hours. The solution was concentrated and collected via vacuum filtration, washed with cold ethyl ether (15 mL) and then dried under vacuum (0.42 g, 41%). $^{1}$H NMR (400 MHz, DMSO) δ 8.80 (s, 1H), 7.98 (ddd, *J* = 8.7, 5.5, 3.2 Hz, 3H), 7.88 – 7.71 (m, 15H), 6.98 (ddd, *J* = 7.7, 1.9, 0.9 Hz, 1H) $^{13}$C NMR (101 MHz, DMSO) δ 167.5, 141.0, 140.9, 136.7, 136.7, 136.5, 136.5, 134.9, 134.8, 132.3, 131.2, 131.0, 127.1, 125.4, 125.3, 123.9, 123.9, 115.2, 114.4, 74.4, 73.8.

**Ethyl (E)-3-(3-oxoisobenzofuran-1(3H)-ylidene)propanoate:**

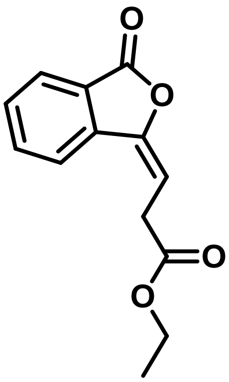

2-((triphenyl-l5-phosphaneylidene)methyl)benzoic acid (2.0 g, 5 mmol, 1 eq) and ethyl 3-oxopropanoate (0.6 g, 5 mmol, 1 eq) were suspended in dichloromethane (15mL) in a 50 mL round bottom flask equipped with a stir-bar. Triethylamine (0.7 mL, 5 mmol, 1 eq) was then added dropwise and the reaction was stirred for 4 hours. The mixture was then diluted with ethyl acetate (25 mL), washed with water and then brine. The organic layers were collected, concentrated and purified by flash chromatography (5% to 20% Hex:EtOAc) to produce a white solid (0.3 g, 21%). *m*/*z* $[M + H]^{+}$ calcd for $C_{13}H_{12}O_4$ 233.1; found 233.6. $^{1}$H NMR (400 MHz, $CDCl_3$) δ 7.95 – 7.85 (m, 1H), 7.77 – 7.63 (m, 2H), 7.62 – 7.45 (m, 1H), 5.31 (t, 1H), 4.27 – 4.09 (m, 2H), 3.54 (d, 2H), 1.24 (t, 3H).$^{13}$C NMR (101 MHz, $CDCl_3$) δ 171.2, 170.1, 146.6, 135.1, 134.1, 129.1, 126.3, 125.8, 122.2, 104.7, 69.8, 31.6, 14.3.

**Ethyl 3-(4-oxo-3,4-dihydrophthalazin-1-yl)propanoate:**

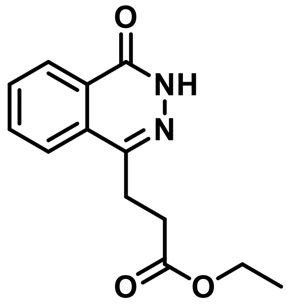

Ethyl (E)-3-(3-oxoisobenzofuran-1(3H)-ylidene)propanoate (0.1 g, 0.41 mmol, 1 eq.) was dissolved in ethanol (15 mL) in a 50 mL round bottom flask equipped with a stir-bar. Hydrazine hydrate (0.04 mL, 0.82 mmol, 2 eq.) was added dropwise to the solution and the reaction was then heated to 50° C for 2 hours. The mixture was allowed to cool to room temperature and the resulting product was collected via vacuum filtration and washed with ethanol (3 x 10 mL) to produce a white solid and was used without further purification (0.2 g, 41%). *m/z* $[M + H]^+$ calcd for $C_{13}H_{14}N_2O_3$ 247.1; found 247.1 $^1$H NMR (400 MHz, $CDCl_3$) δ 8.47 (dt, *J* = 7.7, 1.1 Hz, 1H), 7.88 – 7.84 (m, 2H), 7.78 (ddd, *J* = 8.3, 5.8, 2.6 Hz, 1H), 4.16 (q, *J* = 7.1 Hz, 2H), 3.29 (dd, *J* = 7.8, 6.7 Hz, 2H), 2.86 (dd, *J* = 7.8, 6.7 Hz, 2H), 1.26 (t, *J* = 7.1 Hz, 3H). $^{13}$C NMR (101 MHz, $CDCl_3$) δ 172.9, 160.8, 145.5, 133.7, 131.5, 129.9, 127.7, 127.0, 124.2, 60.7, 30.7, 26.5, 14.2.

**2-chloro-*N*-(3,3-difluorocyclobutyl)acetamide:**

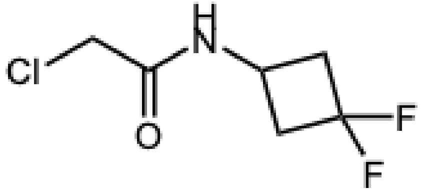

To a 20 mL vial was added a stir bar. This was sealed with a cap and PTFE/rubber septum. The atmosphere of the vial was exchanged by pulling vacuum via a needle inserted into the septum, followed by backfilling with nitrogen. This was done three times total. To this was added 3,3-difluorocyclobutan-1-amine (0.1 mL, 1.05 mmol, 1.05 equiv), $CH_2Cl_2$ (6.6 mL, 0.15 M), and triethylamine (0.28 mL, 2.0 mmol, 2.0 equiv), and the vial was placed into an ice bath and stirred. To this was added, dropwise, chloroacetyl chloride (80 µL, 1.0 mmol, 1.0 equiv). The reaction vessel was then removed from the ice bath and allowed to warm to room temperature. After 4 hours, the reaction was diluted with $CH_2Cl_2$ and transferred to a separatory funnel. The reaction was washed with water, then sat. $NH_4Cl$, then sat $NaHCO_3$, then brine. The organic fraction was then dried over $Na_2SO_4$, filtered, and concentrated. This resulted in a dark blue amorphous solid. This was used without further purification. (149 mg, 81%). $^1$H NMR (400 MHz, $CDCl_3$) δ 6.73 (br s, 1H), 4.30 (ddd, *J* = 6.7, 5.1, 3.2 Hz, 1H), 4.05 (s, 2H), 3.05 (ddt, *J* = 15.5, 11.8, 8.0 Hz, 2H), 2.55 (dddd, *J* = 14.7, 13.1, 11.8, 6.6 Hz, 2H). $^{19}$F NMR (376 MHz, $CDCl_3$) δ -84.97 (d, *J* = 199.7 Hz, 1F), -97.34 (d, *J* = 199.8 Hz, 1F).

**2-chloro-N-(furan-2-ylmethyl)acetamide:**

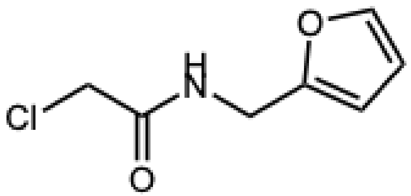

To a 20 mL vial was added a stir bar. This was sealed with a cap and PTFE/rubber septum. The atmosphere of the vial was exchanged by pulling vacuum via a needle inserted into the septum, followed by backfilling with nitrogen. This was done three times total. To this was added furan-2-ylmethanamine (90 µL, 1.05 mmol, 1.05 equiv), $CH_2Cl_2$ (6.6 mL, 0.15 M), and triethylamine (0.28 mL, 2.0 mmol, 2.0 equiv), and the vial was placed into an ice bath and stirred. To this was added, dropwise, chloroacetyl chloride (80 µL, 1.0 mmol, 1.0 equiv). The reaction vessel was then removed from the ice bath and allowed to warm to room temperature. After 4 hours, the reaction was diluted with $CH_2Cl_2$ and transferred to a separatory funnel. The reaction was washed with water, then sat. $NH_4Cl$, then sat $NaHCO_3$, then brine. The organic fraction was then dried over $Na_2SO_4$, filtered, and concentrated. This resulted in a brown amorphous solid. This was used without further purification. (156 mg, 90%). $^1$H NMR (400 MHz, $CDCl_3$) δ 7.38 (d, $J$ = 1.0 Hz, 1H), 6.86 (br s, 1H), 6.34 (dd, $J$ = 3.3, 1.9 Hz, 1H), 6.27 (d, $J$ = 3.0 Hz, 1H), 4.50 (s, 1H), 4.49 (s, 1H), 4.08 (s, 2H).
$^1$H NMR consistent with reported values.

**2-chloro-N-(4-hydroxybenzo[d]oxazol-2-yl)acetamide:**

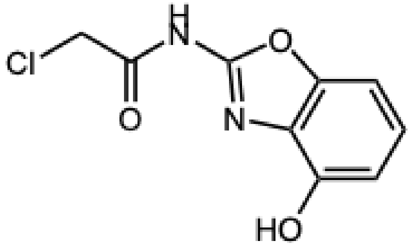

To a 20 mL vial was added 2-aminobenzo[d]oxazol-4-ol (158 mg, 1.05 mmol, 1.05 equiv), and a stir bar. This was sealed with a cap and PTFE/rubber septum. The atmosphere of the vial was exchanged by pulling vacuum via a needle inserted into the septum, followed by backfilling with nitrogen. This was done three times total. To this was added $CH_2Cl_2$ (6.6 mL, 0.15 M), and triethylamine (0.28 mL, 2.0 mmol, 2.0 equiv), and the vial was placed into an ice bath and stirred. To this was added, dropwise, chloroacetyl chloride (80 µL, 1.0 mmol, 1.0 equiv). The reaction vessel was then removed from the ice bath and allowed to warm to room temperature. After 4 hours, the reaction was diluted with $CH_2Cl_2$ and transferred to a separatory funnel. The reaction was washed with water, then sat. $NH_4Cl$, then sat $NaHCO_3$, then brine. The organic fraction was then dried over $Na_2SO_4$, filtered, and concentrated. The crude residue was then purified on using silica gel chromatography (5-40% ethyl acetate in hexanes). This resulted in a white amorphous solid. (51 mg, 22%). **$^1$H NMR** (400 MHz, $CDCl_3$) δ 10.98 (s, 1H), 7.46 (t, *J* = 8.4 Hz, 1H), 7.07 (dd, *J* = 8.3, 0.9 Hz, 1H), 7.04 (dd, *J* = 8.5, 0.9 Hz, 1H), 4.70 (s, 2H).

**2-(2-chloroacetamido)-4,5,6,7-tetrahydrobenzo[b]thiophene-3-carboxamide:**

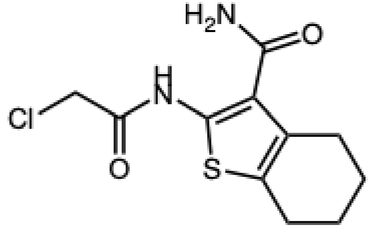


To a 20 mL vial was added 2-amino-4,5,6,7-tetrahydrobenzo[b]thiophene-3-carboxamide (206 mg, 1.05 mmol, 1.05 equiv), and a stir bar. This was sealed with a cap and PTFE/rubber septum. The atmosphere of the vial was exchanged by pulling vacuum via a needle inserted into the septum, followed by backfilling with nitrogen. This was done three times total. To this was added $CH_2Cl_2$ (6.6 mL, 0.15 M), and triethylamine (0.28 mL, 2.0 mmol, 2.0 equiv), and the vial was placed into an ice bath and stirred. To this was added, dropwise, chloroacetyl chloride (80 µL, 1.0 mmol, 1.0 equiv). The reaction vessel was then removed from the ice bath and allowed to warm to room temperature. After 4 hours, the reaction was diluted with $CH_2Cl_2$ and transferred to a separatory funnel. The reaction was washed with water, then sat. $NH_4Cl$, then sat $NaHCO_3$, then brine. The product was found as a solid suspended in the aqueous layer. The solid was then suspended in $CH_2Cl_2$ and filtered through a fine glass frit. This was washed with $CH_2Cl_2$, and dried over vacuum, resulting in a dusty brown solid. (192 mg, 70%). **$^1$H NMR** (400 MHz, DMSO) δ 12.24 (s, 1H), 7.82 – 6.94 (m, 4H), 4.50 (s, 2H), 2.70 (d, *J* = 4.9 Hz, 2H), 2.68 – 2.60 (m, 2H).

**3-(3-(2-((3,3-difluorocyclobutyl)amino)-2-oxoethyl)-4-oxo-3,4-dihydrophthalazin-1-yl)propanoic acid (Compound 13):**

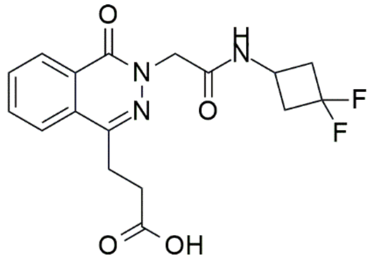


Potassium iodide (10 mg, 0.06 mmol, 1.5 eq), 2-chloro-N-(3,3-difluorocyclobutyl)acetamide (11 mg, 0.06 mmol, 4 eq), and acetone (4 mL) were combined in a round bottom flask equipped with a reflux condenser and stir-bar. The reaction stirred for 30 minutes before potassium carbonate (8.3 mg, 0.06 mmol, 1.5 eq) and ethyl 2-(4-oxo-3,4-dihydrophthalazin-1-yl)propanoate (10 mg, 0.04 mmol, 1 eq.) in acetone (6 mL) were added. The mixture was then heated to reflux for 4 hours. After cooling to room temperature, the solution was diluted with water (25 mL) and extracted using ethyl acetate (3 x 25 mL). The crude product was then added to a solution of THF/MeOH (8 mL, 1:1), lithium hydroxide (1 mL, 1M), and stirred overnight. The reaction was then diluted with water and titrated using hydrochloric acid (1M) until a pH below 3 was attained, resulting in an opaque white solid. The compound was then collected vacuum filtration and washed with water and further purified using semi-preparative HPLC to afford Compound 13 as an opaque white solid (1.7 mg, 11%). HRMS $[M + Na]^+$ calcd for $C_{17}H_{17}F_2N_3O_4$ 383.1084; found 383.1146; Δppm = 3.96. $^1$H NMR (400 MHz, MeOD) δ 8.39 (d, *J* = 8.0 Hz, 1H), 8.13 (d, *J* = 8.2 Hz, 1H), 7.97 (t, *J* = 7.6 Hz, 1H), 7.87 (t, *J* = 7.8 Hz, 1H), 4.25-4.19 (m, 2H), 3.50 (s, 2H), 2.91 (s, 2H), 2.67 (t, *J* = 7.7 Hz, 1H).

**3-(3-(2-((furan-2-ylmethyl)amino)-2-oxoethyl)-4-oxo-3,4-dihydrophthalazin-1-yl)propanoic (Compound 14):**

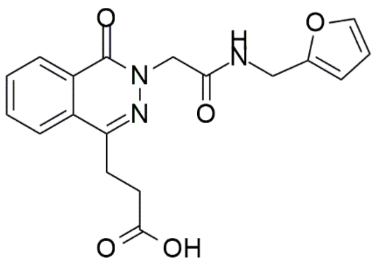


Potassium iodide (10 mg, 0.06 mmol, 1.5 eq), 2-chloro-N-(furan-2-ylmethyl)acetamide (6.8 mg, 0.06 mmol, 1.5 eq), and acetone (4 mL) were combined in a round bottom flask equipped with a reflux condenser and stir-bar. The reaction stirred for 30 minutes before potassium carbonate (8.3 mg, 0.06 mmol, 1.5 eq) and ethyl 2-(4-oxo-3,4-dihydrophthalazin-1-yl)propanoate (10 mg, 0.04 mmol, 1 eq) in acetone (6 mL) were added. The mixture was then heated to reflux for 4 hours. After cooling to room temperature, the solution was diluted with water (25 mL) and extracted using ethyl acetate (3 x 25 mL). The crude product was then added to a solution of THF/MeOH (8 mL, 1:1), lithium hydroxide (1 mL, 1M), and stirred overnight. The reaction was then diluted with water and titrated using hydrochloric acid (1M) until a pH below 3 was attained, resulting in an opaque white solid. The compound was then collected vacuum filtration and washed with water and further purified using semi-preparative HPLC to afford Compound 14 as an opaque white solid (4.9 mg, 34%). HRMS $[M + H]^+$ calcd for $C_{18}H_{17}N_3O_5$ 378.1065; found 378.1048; Δppm = -3.34. $^1$H NMR (400 MHz, MeOD) δ 7.72 – 7.63 (m, 2H), 7.58 (m, 2H), 7.45 – 7.41 (m, 1H), 6.36 (dd, J = 3.3, 1.8 Hz, 1H), 6.26 (d, J = 3.3 Hz, 1H), 4.43 (s, 2H), 3.92 (s, 2H), 1.35 – 1.29 (m, 2H), 1.26 - 1.23 (m, 2H). $^{13}$C NMR (101 MHz, MeOD) δ 170.98, 151.54, 141.89, 132.40, 132.37, 131.73, 131.64, 128.65, 128.52, 127.39, 126.64, 109.94, 106.71, 71.21, 58.11, 35.14, 29.37, 28.45.

**3-(3-(2-((3-carbamoyl-4,5,6,7-tetrahydrobenzo[b]thiophen-2-yl)amino)-2-oxoethyl)-4-oxo-3,4-dihydrophthalazin-1-yl)propanoic acid (Compound 15):**

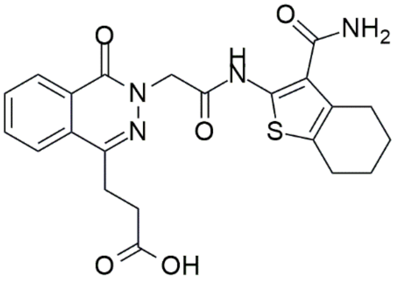


Potassium iodide (10 mg, 0.06 mmol, 1.5 eq), 2-(2-chloroacetamido)-4,5,6,7-tetrahydrobenzo[b]thiophene-3-carboxamide (16 mg, 0.06 mmol, 1.5 eq), and acetone (4 mL) were combined in a round bottom flask equipped with a reflux condenser and stir-bar. The reaction stirred for 30 minutes before potassium carbonate (8.3 mg, 0.06 mmol, 1.5 eq) and ethyl 2-(4-oxo-3,4-dihydrophthalazin-1-yl)propanoate (10 mg, 0.04 mmol, 1 eq) in acetone (6 mL) were added. The mixture was then heated to reflux for 4 hours. After cooling to room temperature, the solution was diluted with water (25 mL) and extracted using ethyl acetate (3 x 25 mL). The crude product was then added to a solution of THF/MeOH (8 mL, 1:1), lithium hydroxide (1 mL, 1M), and stirred overnight. The reaction was then diluted with water and titrated using hydrochloric acid (1M) until a pH below 3 was attained, resulting in an opaque white solid. The compound was then collected vacuum filtration and washed with water and further purified using semi-preparative HPLC to afford Compound 15 as an opaque white solid (2.2 mg, 12%). HRMS $[M + H]^+$ calcd for $C_{22}H_{22}N_4O_5S$ 455.1389; found 455.1387; Δppm = 0.83 $[M + Na]^+$ calcd 477.1; found 477.2. $^1H$ NMR (400 MHz, MeOD) δ 8.00 (dd, $J$ = 7.9, 1.4 Hz, 1H), 7.67 (ddd, $J$ = 7.0, 4.1, 1.5 Hz, 2H), 7.60 – 7.55 (m, 1H), 7.44 – 7.35 (m, 1H), 4.42 (d, $J$ = 2.3 Hz, 1H), 3.42 (s, 1H), 3.01 – 2.94 (m, 1H), 2.87 (td, $J$ = 7.0, 4.3 Hz, 1H), 2.83 – 2.76 (m, 1H), 1.95-1.81 (m, 2H) 1.32 (d, $J$ = 7.5 Hz, 1H). $^{13}C$ NMR (101 MHz, MeOD) δ 134.07, 132.38, 132.12, 131.73, 131.63, 130.59, 128.69, 128.53, 127.42, 126.66, 122.31, 109.95, 71.20, 70.05, 62.38, 58.12, 57.98, 35.14, 29.35, 25.23, 22.63.

**Molecular Docking:** Conformers used in this study were generated using the OEOMEGA tool from the OpenEye suite. They were also modified to adopt the proper protonation state at a pH of 7.4 using the FILTER tool. Docking experiments were performed with FRED (OpenEye) and docking poses and scores were then analyzed manually in VIDA (OpenEye).

**Liquid Chromatography-Mass Spectrometry (LC-MS):** Liquid chromatography-mass spectrometry was performed on inhibitors following chemical synthesis (Agilent 1260 Infinity II/Infinity Lab LC/MSD XT). Samples dissolved in variable DMSO concentrations were run on the LC-MS using a 10 minute 5-95% gradient of acetonitrile in water with a formic acid modifier (0.1%).

**Protein Expression:** His-tag HDAC6$^{1109-1215}$ was first raised from bacterial glycerol stock containing HDAC6 plasmid (Addgene:3gv4) in DH5α competent E. coli cells. Bacteria were streaked on a kanamycin-treated agar plate and given 24 hours to grow. A single colony was then selected and transferred to a starter culture containing 1:1000 dilution of Kanamaycin (50 mg/mL stock) in Luria-Betani (LB) broth incubated for 24 hours. The QIA Mini Prep kit procedure was used the following day to collect plasmid from DH5a cells. 5 µL of isolated plasmid was then transformed into Rosetta2 competent cells and incubated together on ice for 30 min. Following incubation, the treated cells were heat shocked at 42 °C for 45 sec. 400 µL of SOC media was added to the solution and incubated for 1 hour at 37 °C. After the initial incubation period, 100 µL of cells in SOC media were transferred to kanamycin plates and permitted to grow for 18-24 hr.

Following this incubation, 1 L and 100 mL quantities LB broth were prepared with 20-25 g and 2.0-2.5 g respectively from Fisher Bioreagents dry LB powder in Erlenmeyer flasks. LB was then autoclaved and permitted to cool. 1:1000 dilution of Kanamycin was added to both flasks. 50 µL of $ZnSO_4$ was added only to the 1 L flask. Colonies from the Rosetta competent cells were selected and added to the 100 mL flask to grow up on a shaker for 18-24 hours at 37 °C. The following day, 10-15 mL of the 100 mL liquid culture was transferred to the 1 L Erlenmeyer flask. The 1 L flask was incubated on a shaker for 2-3 hours at 37 °C. Following this incubation period, OD600 absorbance data was collected using a Nanodrop. Indication of an OD600 between 0.4-0.8 demonstrated sufficient growth of the cells. The 1 L flask was then induced with 200 µM of IPTG and incubated overnight (18-24 hours) at 16 °C.

**Protein Purification:**

***Bacterial Lysis:*** Following protein expression, liquid cultures were divided into 250 mL Nalgene centrifuge bottles and spun down at 6500 rpm at 4 °C for 20 minutes in the Sorvall Lynx 4000 Centrifuge (Thermo Scientific). After initial spin down, the LB broth was decanted leaving behind pelleted His-tag $HDAC6^{1109\text{-}1215}$ containing bacterium. Pellets were resuspended in 7 mL of fresh LB brother and transferred to 50 mL Nalgene centrifuge bottles and spun down at 4500 rpm at 4 °C for 20 minutes. The pellet was resuspended in 20-25 mL of lysis buffer (2 mM DTT, 5 mM imidazole, a protease inhibitor tablet (Pierce), Base Buffer [50 mM Tris (pH 8.0), 300 mM NaCl]) and sonicated for 10 minutes (40% amplitude, 2 seconds on/off). Following sonication, lysate was spun down at 13,000 rpm at 4 °C for 60 minutes and added to a nickel column.

***Nickel Column Purification:*** Nickel resin beads (Qiagen Ni-NTA Agarose) were added and washed twice on a column with 25 mL of cold MilliQ water. 25-30 mL of lysis buffer and clarified lysates were then added to the beads on the column and placed in the cold room on a rocker for 2-24 hr. Flowthrough from the column was collected in a 50 mL canonical tube. The column was then washed 2x with 25 mL Wash Buffer (2 mM DTT, 50 mM imidazole, Base Buffer [50 mM Tris (pH 8.0), 300 mM NaCl]). Flow through from each was collected in 50 mL canonical tubes. Following washing steps, His-tag $HDAC6^{1109\text{-}1215}$ along the resin beads was eluted using Elution Buffer (2 mM DTT, 250 mM imidazole, Base Buffer [50 mM Tris (pH 8.0), 300 mM NaCl) 3 times. Eluant was collected in 15 mL canonicals after each round of elution buffer treatment.

***SDS-Page:*** 7 µL of lysate flowthrough, wash buffer elution, and elution buffer elution and 3 µL of 2-mercaptoethanol treated 4X Sample Buffer (BioRad) were combined in 1.5 ml microcentrifuge tubes. Each tube was heated for 3-5 minutes at 95 °C on a heat block and then loaded into an SDS-page gel (Miniprotean TGX gels, 4-20%, 15 well, 15 μL/well). The gel was run for 35 minutes at 180 V and then transferred to a gel box and stained with Coomassie. Coomassie imaging was done on the Molecular Imager ChemiDoc XPS + imaging system (BioRad). Band detection for the presence of His-tag HDAC6$^{1109-1215}$ at 13-14 kDa was performed using the Image Lab 6.0.1 software. Coomassie gels were run similarly on His-tag cleaved HDAC6$^{1109-1215}$. Positive results indicated a 1 kDa downward shift of the HDAC6$^{1109-1215}$ along the protein gel.

***Dialysis Buffer Exchange and Protein Concentration***: 1 L of Base Buffer [50 mM Tris (pH 8.0), 300 mM NaCl] was prepared then a dialysis buffer exchange system was set up using 7,000 MWCO dialysis tubing (Thermo Scientific Snakeskin Dialysis Tubing). Purified His-tag HDAC6$^{1109-1215}$ was added to the snakeskin and then into the base buffer container with a spin bar and left to exchange for 12-24 hours. Purified his-tag HDAC6$^{1109-1215}$ was collected the following day and added to protein concentration tubes (Sigma Aldrich Millipole centrifugal filter units) with 3,000 MWCO cut off and spun down for 10 minutes intervals at 4500 rpm, 4 °C. The molar extinction coefficient for HDAC6$^{1109-1215}$ was calculated through the Expasy Protparam and the protein concentration was measured at A280 using a Nanodrop. Following protein measurement, purified His-tag HDAC6$^{1109-1215}$ was added to 1.5 mL microcentrifuge vials.

***Thrombin Cleavage:*** Thrombin CleanCleave kit thrombin beads (Sigma Aldrich) were resuspended in thrombin-agarose resin to produce a homogeneous mixture and prepared using the CleanCleave kit procedure. Beads were then added to the purified His-tag HDAC6$^{1109-1215}$ solution and left to rock for 12-20 hours at room temperature (RT). Beads were spun down at 4°C, 500 x rcf the following day and the supernatant was decanted into fresh 1.5 mL microcentrifuge tubes. A Coomassie was run on the cleaved HDAC6$^{1109-1215}$ and protein concentrations were measured on the Nanodrop. Fully cleaved HDAC6$^{1109-1215}$ was flash-frozen using liquid nitrogen and stored at -20 °C.

***Reverse Nickel:*** Reverse nickel was performed on any partially cleaved His-tag HDAC6$^{1109-1215}$ / HDAC6$^{1109-1215}$ purified proteins. Nickel column was set up using nickel resin beads and purified protein was added to the column. Flowthrough containing purified cleaved HDAC6$^{1109-1215}$ was collected in 15 mL canonical tubes and was washed 2x with 2 mL of Wash Buffer and 3x with 2 mL of Elution Buffer. Coomassie gel was run following reverse nickel column separation and purified his-tag HDAC6 $^{1109-1215}$ / purified cleave HDAC6 $^{1109-1215}$ were found via protein band identification. Purified protein was concentrated and measured using the Nanodrop, then flash-frozen using liquid nitrogen and stored at -20 °C.

***Superdex 75 (S75) Size Exclusion Chromatography:*** HDAC6$^{1109-1215}$ purification was performed using the Superdex 75 size exclusion column on an AKTA Pure chromatography system. The column was equilibrated overnight using 50 mM Tris (pH 8.0), 300 mM NaCl solution. Following equilibration, concentrated (1%-5% of CV) HDAC6$^{1109-1215}$ in 50 mM Tris (pH 8.0), 300 mM NaCl buffer was injected into the column with 1-2 mL fractions collected of peaks. Peak fractions were collected following size separation and run on SDS page gels then visualized with Coomassie staining (Fig. S1)

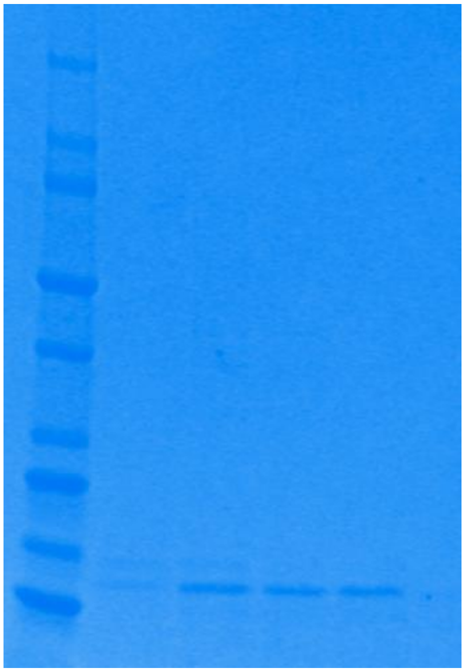

**Figure S1 – SDS-PAGE for Purified HDAC6$^{109\text{-}1215}$ ZnF UBD**

**Fluorescence polarization competition assays:**

FPA experiments were conducted in 384-well black polystyrene plates (Fisher). Assay buffer, HDAC6$^{1109\text{-}1215}$, FITC-RLRGG peptide tracer (Genescript), and the ZnF-UBD ligands were added to the first 10 treatment wells with the 11th well containing the control treatment (assay buffer + FITC-RLRGG peptide). Assay buffer was comprised of 10 mM of HEPES, 150 mM of NaCl, and 10 mg of BSA in 10 mL of MilliQ water solvent. HDAC6$^{1109\text{-}1215}$, FITC-RLRGG tracer and ZnF-UBD ligands were diluted in dilution buffer containing 10 mM of HEPES, 150 mM of NaCl, brought up to 10 mL in MilliQ water prior to 384 well plate addition. FITC-RLRGG peptide was serially diluted from an initial 1 mM stock solution to a 10 µM solution, to a final concentration of 160 nM. 12.5 µL of 160 nM FITC-RLRGG peptide was added to each treatment well in 384 well plates. Purified HDAC6$^{1109\text{-}1215}$ was diluted from a purification-dependent initial stock concentration to 10 µM. 12.5 µL of the 10 µM stock was then added to the 384 well plate. ZnF-UBD ligands were similarly subjected to a dilution series prior to FPA analysis. 10 dilutions series were performed in a 96-well black polystyrene flat bottom plates (Fisherbrand) with concentrations ranging from 800 µM to 0.0031 µM. Dilutants from each well were then added to the 384 well plate, alongside the FITC-LRLGG tracer, assay buffer, and HDAC6$^{1109\text{-}1215}$ purified protein. Following reagent addition, the plate was then spun for 40 sec at 1200 rpm. Following a 1 hr incubation period, FITC fluorescence polarization was measured on a plate reader (Tecan

Spark) with an excitation wavelength of 480 nm and an emission wavelength of 528 nm. Using GraphPad Prism, anisotropic values were imported using dose-response where X is log(dose) and normalized (average sub-column and normalize means, smallest value = 0, largest value = 100, report data as percentages, bottom constraint = 0). Normalized data was plotted using a nonlinear regression (curve fit).

**Differential Scanning Fluorimetry (DSF):** DSF was performed in a 100 µL reaction mixture and completed in the succeeding order. 50 µL of 50 mM Tris (pH 8.0), 300 mM NaCl buffer was added to a 1.5 mL Eppendorf tube, followed by MilliQ water. Purified HDAC6$^{1109\text{-}1215}$ was concentrated to a final concentration of 1.0 mg/mL via Nanodrop. 10 µM of purified HDAC6$^{1109\text{-}1215}$ was added to an experimental Eppendorf. MilliQ volume addition was adjusted according to the 10 µM HDAC6$^{1109\text{-}1215}$ only conditions. DMSO stocks of compounds were diluted to 10 mM and added into the 100 µL reaction for a final concentration of 50 µM. Following compound addition, 5000X SYPRO Orange Dye (Invitrogen) was diluted to a 100X concentration in 50 mM Tris (pH 8.0) and 300 mM NaCl buffer, and 10 µL was added into the 100 µL reaction. Buffer only controls contained equal volumes of MilliQ (50 µL) and 50 mM Tris (pH 8.0), 300 mM NaCl buffer (50 µL). Buffer + dye controls contained MilliQ (40 µL), 50 mM Tris (pH 8.0), 300 mM NaCl buffer (50 µL), and 100X SYPRO Orange Dye (10 µL). Reaction master mixes were incubated on ice for 20-30 minutes. Following incubation, 3 replicates of 20 µL of each eaction mix were pipetted into MicroAmp Optical 96-Well Reaction Plates (ThermoFisher) and spun down for 3 minutes at 300 rpm. DSF was run on Quant Studio 3 (Thermo Fisher) using the QuantStudio Design and Analysis Software v1.5.3. Experimental results were plotted using GraphPad prism. First derivative plots were calculated through the XY analysis smooth, differentiate or integrate curve function. Averaged fluorescence values were plotted using a standard XY graph (Y= Temperature).

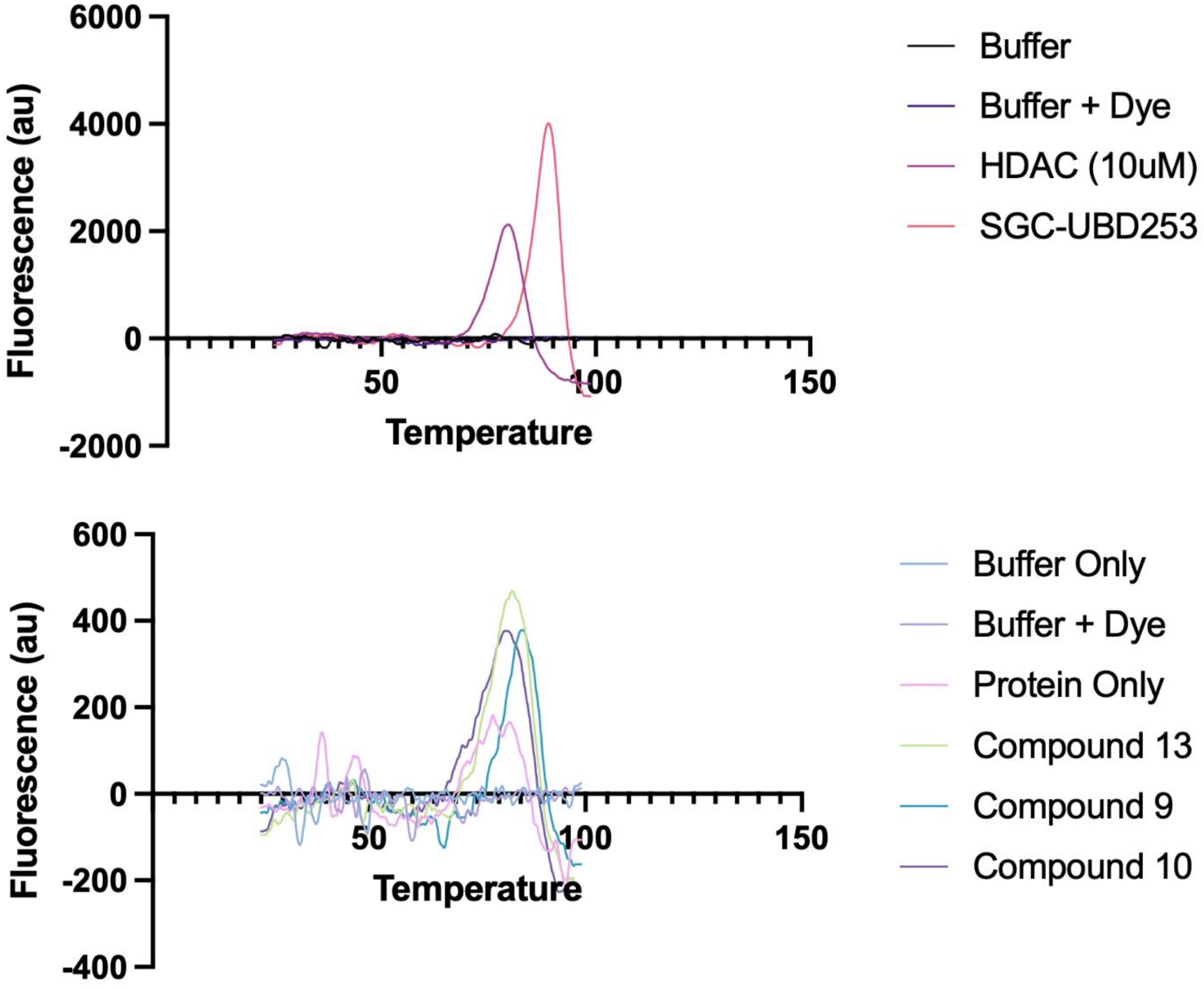


Figure. S2 - First Derivatives of DSF Curves

Table S1. Calculated Molecular Properties

| Cmpd | ClogP | MW | # HBA | # HBD | Fraction SP$^3$ | TPSA | # Rot. Bonds | # Ar. Rings | LOSERS |
|---|---|---|---|---|---|---|---|---|---|
| **7** | 3.08 | 452.15 | 8 | 4 | 0.30 | 127.31 | 8 | 3 | 38.71 |
| **8** | 1.90 | 408.11 | 10 | 3 | 0.15 | 147.55 | 6 | 4 | 42.60 |
| **9** | 1.32 | 355.12 | 8 | 2 | 0.22 | 114.43 | 7 | 3 | 57.54 |
| **10** | 1.33 | 365.12 | 7 | 2 | 0.41 | 101.29 | 6 | 2 | 73.55 |

**Individual Competition Assays:**

SGC-UBD 253

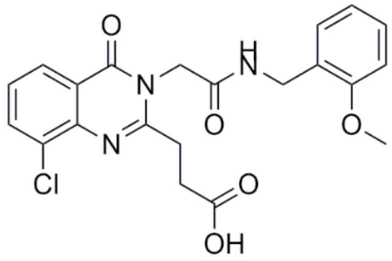

Molecular Weight: 429.857

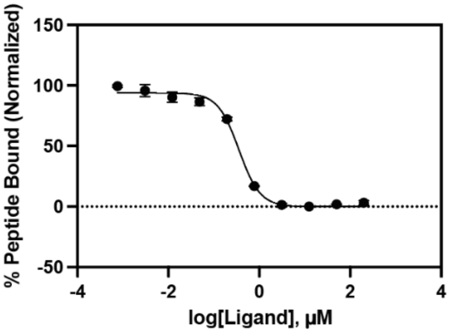


Compound 5

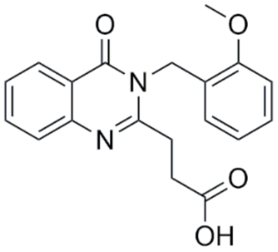

Molecular Weight: 338.363

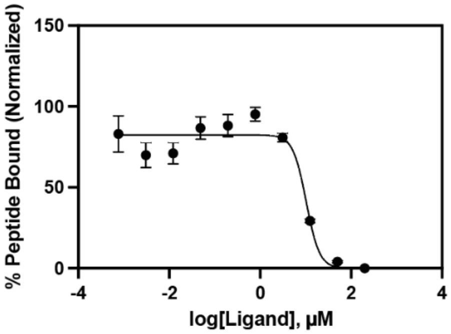


Compound 6

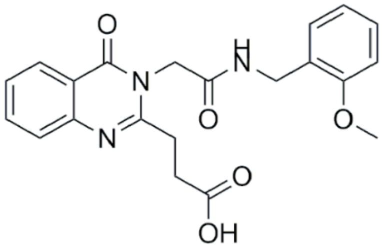

Molecular Weight: 395.415

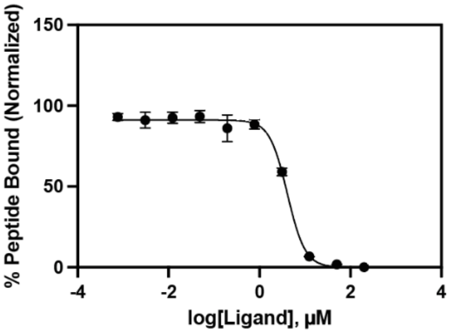

Compound 7

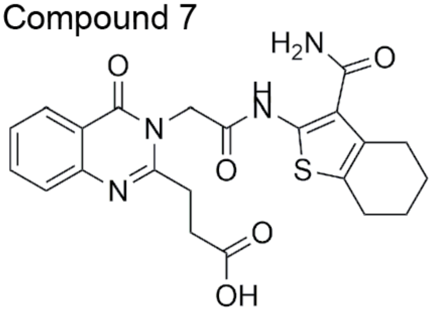


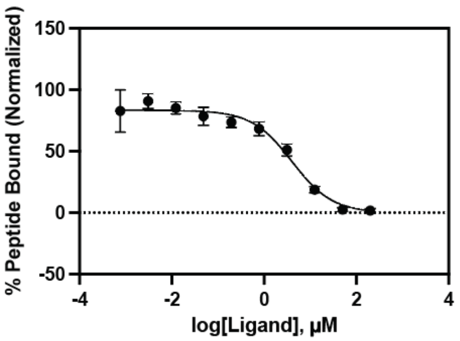


Molecular Weight: 454.501

Compound 8

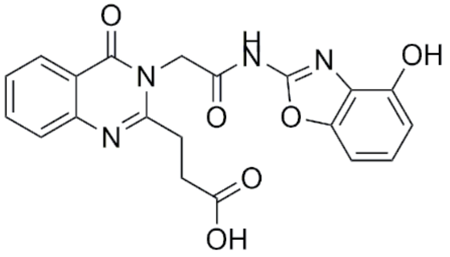


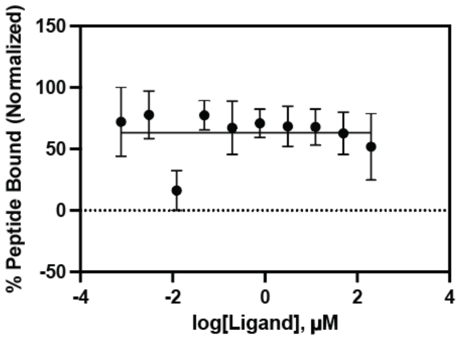


Molecular Weight: 408.370

Compound 9

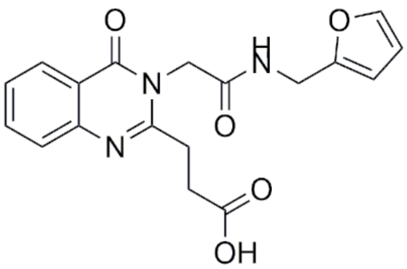


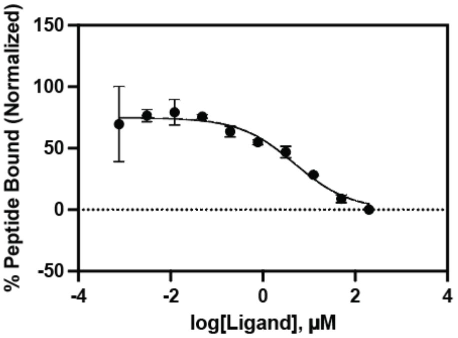


Molecular Weight: 355.350

Compound 10

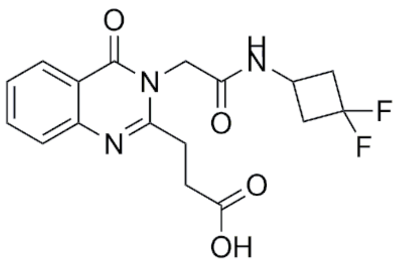


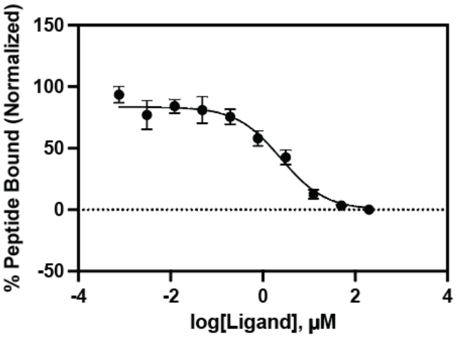


Molecular Weight: 365.337

Compound 11

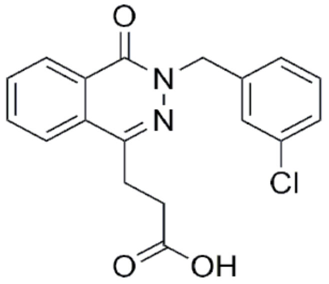

Molecular Weight: 342.779

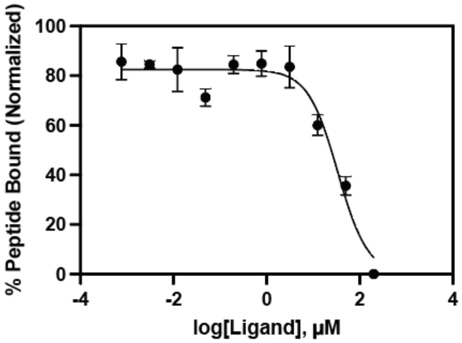


Compound 12

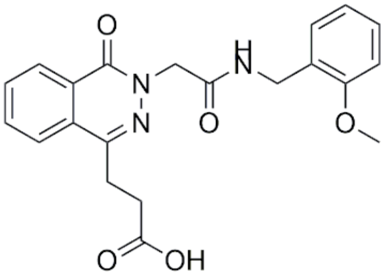

Molecular Weight: 395.415

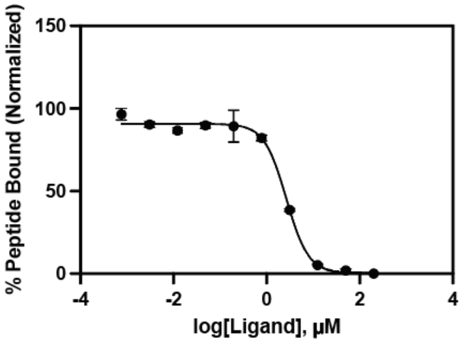


Compound 13

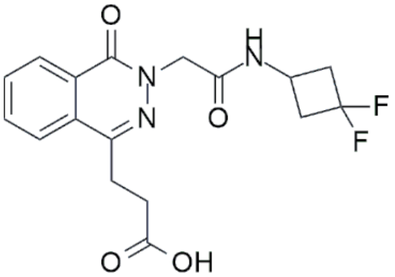

Molecular Weight: 365.33

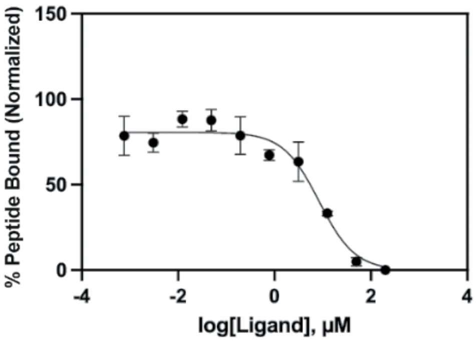


Compound 14

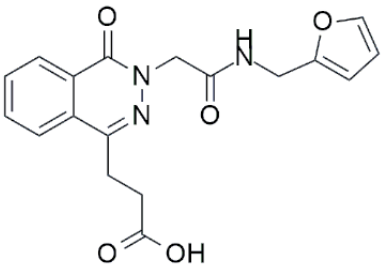

Molecular Weight: 355.34

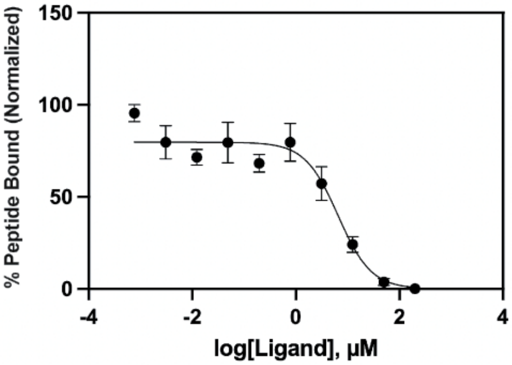

Compound 15

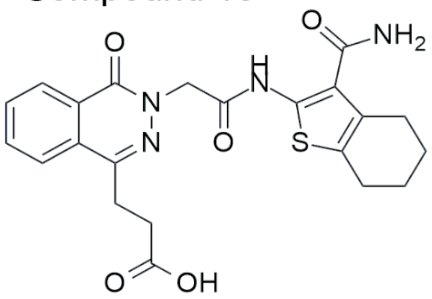


Molecular Weight: 454.50

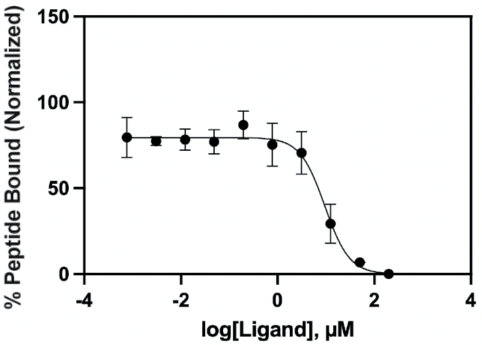
